# AP2-L-Dependent Processes Promote PVM Maintenance and *Plasmodium* Liver-stage Development

**DOI:** 10.64898/2026.10.29.762720

**Authors:** Destaalem T. Kidane, Anusha Thaniana, Farhana Mosaddeque, Camille Louvat, Amit Subudhi, Arnab Pain, Purnima Bhanot

## Abstract

Successful malaria infection requires transformation of *Plasmodium* sporozoites into replicative exo-erythrocytic forms (EEFs) within hepatocytes, a process requiring a functional parasitophorous vacuole membrane (PVM) and interactions with host cellular pathways. ApiAP2 transcription factors regulate developmental progression throughout the *Plasmodium* life cycle. AP2-L consists of an extended amino terminal region and two AP2 domains and is highly expressed in asexual stage trophozoites, sporozoites and early EEFs, but its role in these stages is poorly understood. We characterized the functional consequences of deleting the entire *P. berghei* ap2-l coding sequence on the erythrocytic cycle *in vivo*, sporozoite gene expression, liver-stage development, and hepatocyte-parasite interactions. AP2L-KO parasites maintained erythrocytic replication, but sporozoite inoculation failed to establish blood-stage infection in mice. AP2-L loss decreased transcripts encoding proteins involved in sporozoite gliding motility, host-cell entry, and rhoptry secretion. Motif discovery upstream of a subset of these genes suggests possible direct targets of AP2-L. Infected HepG2 cells showed reduced EEF numbers and sizes, impaired parasite DNA replication, and abnormal parasite positioning relative to the host nucleus. PVMs of AP2L-KO parasites exhibited aberrant localization of proteins, absence of tubular vacuolar network extensions, and sensitivity to detergent. These abnormalities were accompanied by increased accumulation of the host’s autophagy-associated protein around parasites, suggesting efficient host-mediated elimination. Additionally, AP2L-KO parasites displayed reduced association with the host microtubule-organizing center, Golgi compartments, and COPB2-positive vesicular structures, suggesting an impairment in uptake of nutrients. Together, these findings demonstrate that AP2-L directs both vacuolar biogenesis and host cytoskeletal/vesicular capture in liver-stage parasites.

**Importance:** Before causing symptomatic malaria, *Plasmodium* parasites multiply silently inside liver cells within a specialized protective structure called the parasitophorous vacuole. Understanding how the parasite builds and defends this essential niche is critical for developing strategies to prevent malaria. In this study, we demonstrate that the parasite transcription factor AP2-L controls a gene expression program vital for liver-stage survival. By analyzing parasites with a complete gene deletion, we show that AP2-L, directly or indirectly, activates genes required for vacuole membrane formation and maintenance. Without AP2-L, parasites form compromised vacuoles, fail to evade host cell autophagic clearance, and cannot capture host cell organelles needed for growth, completely halting infection. These findings establish AP2-L as a key regulator connecting nuclear gene expression to vacuole integrity and host cell manipulation, processes that are fundamental to intracellular pathogen survival.

## Introduction

*Plasmodium* parasites undergo a complex life cycle that alternates between an *Anopheles* mosquito vector and a vertebrate host. Following transmission via a female *Anopheles* mosquito bite, sporozoites enter the bloodstream, migrate towards the liver and invade hepatocytes, where they establish the clinically silent but obligatory liver-stage (LS) of infection. Within hepatocytes, elongate sporozoites remodel into spherical exo-erythrocytic forms (EEFs), preferentially in the juxtanuclear region of the hepatocyte (Bano et al., 2007) (Spielmann et al., 2012), undergoing repeated rounds of closed mitosis (Gerald et al., 2011) (Jayabalasingham et al., 2010). This rapid proliferative phase generates thousands of merozoites which are subsequently released into the bloodstream to initiate the erythrocytic stage of infection responsible for malaria pathology (Prudêncio et al., 2006) (Sturm et al., 2009). Although the transformation from sporozoite to an EEF can occur in the absence of host cells, productive growth, nuclear replication and differentiation into merozoites require intracellular development within hepatocytes (Kaiser et al., 2003).

Productive hepatocyte invasion is accompanied by formation of a parasitophorous vacuole (PV), which separates the developing parasite from the host-cell cytoplasm. The surrounding membrane that delimits the PV, the PV membrane (PVM) constitutes the major interface between parasite and host. The PVM is derived from the hepatocyte plasma membrane and is rapidly modified by parasite-derived proteins released from rhoptries (Ghosh et al., 2017) during invasion, creating a specialized membrane compartment with functions to acquire nutrients for the growing parasite, dispose its waste products and protect it against the intracellular defenses of the hepatocyte (Spielmann et al., 2012). The PVM facilitates exchange with the host cytosol by allowing small molecules to enter the PV, supporting the metabolic demands of the rapidly growing parasite. In addition, specific PVM proteins mediate interactions with host proteins to obtain nutrients. For example, UIS3 interacts with host liver-fatty acid binding protein (L-FABP) and EXP1 interacts with apolipoprotein H (ApoH) (Sa et al., 2017) (Mikolajczak et al., 2007).

The host-parasite interface extends beyond the PVM. Membranous extensions originating from the PVM form the dynamic tubulovesicular network (TVN), which projects into the host-cell cytoplasm in the form of membrane clusters, vesicles and thin tubules (Nyboer et al., 2018) (Grutzke et al., 2014). The PVM and TVN interact with several components of the host’s endomembrane system, including late endosomes, lysosomes, and the Golgi complex (Nyboer et al., 2018) (Grutzke et al., 2014). Components of the host autophagy related machinery are also recruited to the parasite-associated membranes (Agop-Nersesian et al., 2017) (Prado et al., 2015) (Thieleke-Matos et al., 2016). Soon after invasion, microtubule-associated protein 1 light chain 3 (LC3), a central protein in the autophagy pathway, is recruited to the PVM. Persistent LC3 association is associated with impaired parasite growth and elimination via an autophagy-like mechanism (Prado et al., 2015). Surviving parasites, however, progressively reduce LC3 accumulation around the parasite body by transporting it into the TVN and eventually shedding it into the cytosol, highlighting the parasite’s ability to regulate this potentially detrimental host response (Prado et al., 2015) (Agop-Nersesian et al., 2017). Thus, the PVM functions as a dynamic interface that supports nutrient acquisition, host-cell exploitation, and protection from intracellular defenses.

The extensive cellular remodeling that accompanies the transition from sporozoite to a replicating EEF requires coordinated changes in parasite gene expression. A major family of transcriptional regulators in *Plasmodium* is the apicomplexan ApiAP2 family, whose members contain one or more Apetala2 (AP2) DNA-binding domains (Painter et al., 2011). Genetic studies have established essential functions for ApiAP2 proteins during different developmental transitions, including gametocytogenesis, ookinete development, sporozoite formation and LS maturation (Kafsack et al., 2014) (Yuda et al., 2010) (Yuda et al., 2009) (Iwanaga et al., 2012). Several ApiAP2 proteins display sequence specific DNA binding preferences, with AP2 domains recognizing distinct motifs, supporting their role in coordinating stage specific gene expression throughout the parasite life cycle (Campbell et al., 2010). However, how transcriptional regulation coordinates the developmental remodeling and host interactions required for productive hepatocyte infection remains poorly understood.

AP2-L expression is detected in human-infective *P. falciparum* and *P. vivax* sporozoites (Zanghi et al., 2018) (Muller et al., 2019). In the rodent-infective *P. berghei,* AP2-L protein localizes to the nuclei of blood stage trophozoites and salivary gland (SG) sporozoites (Iwanaga et al., 2012). During LS development, fluorescently tagged AP2-L protein is detectable by 12 h p.i., increases in abundance during schizogony and disappears upon formation of mature liver schizonts (Iwanaga et al., 2012). AP2-L contains two closely spaced AP2 DNA-binding domains near its C-terminus followed by an additional highly conserved ACDC (AP2-coincident-domain mainly at the C-terminus) domain. Attempts to delete the ap2-l orthologs in *P. yoelii* (PY17X_021580) and *P. falciparum* (PF3D7_0730300) were unsuccessful (Zhang et al., 2017) (Shang et al., 2021) but a genome-wide piggyBac saturation mutagenesis screen in *P. falciparum* recovered an insertion between the two AP2 domains of PF3D7_0730300 (Zhang et al., 2018). In *P. berghei*, a previous genetic disruption deleted the C-terminal region of ap2-l gene (PBANKA_0214400), including both AP2 domains and the ACDC domain, while leaving intact the proximal two-thirds of the gene that constitutes a substantial portion of the N-terminal region (Iwanaga et al., 2012). It is possible that the *P. falciparum* transposon mutant and the *P. berghei* mutant parasites make truncated AP2L, leaving open the question of AP2-L’s function in the parasite.

Here, we investigated the function of AP2-L using a complete deletion of the *P. berghei* ap2-l coding sequence, combined with *in vivo* infection studies, quantitative analyses of LS development and transcriptomic profiling of SG sporozoites. We show that AP2-L regulated processes are essential for the formation of a robust PVM, nuclear replication in the hepatic trophozoites, evasion of host autophagy-like pathways, and nutrient uptake via hijacked vesicular traffic. Transcriptomic analyses further reveal broad dysregulation of genes involved in host-cell interaction, motility, metabolism, and other processes required for successful infection. Together, these findings identify AP2-L as a major regulator of a sporozoite transcriptional program that enables the parasite to establish and maintain the intracellular niche required for productive LS development.

## Results

### Predicted structure of AP2-L

AP2-L proteins contain two AP2 domains and an adjacent ACDC domain at their carboxy termini, and an extended amino terminal region (Iwanaga et al., 2012). Multiple sequence alignment of AP2-L orthologs revealed substantially greater conservation within the carboxy-terminal region than across the extended amino terminus, which contained localized conserved segments interspersed among highly variable regions and extensive alignment gaps. Alignment of the two AP2 domains from 21 *Plasmodium* orthologs revealed high amino acid conservation in both domains, with the first AP2 domain exhibiting greater conservation (98 % mean identity) than the second (79.5 % mean identity) (**Supplementary Fig. 1A)**. Structural predictions of the amino terminal regions of *P. berghei and P. falciparum*, using AlphaFold3, reveal largely unstructured regions with a moderate possibility of alpha helices and beta strands **(Supplementary Fig. 1B)**. We analyzed the *P. berghei and P. falciparum* sequence using Rapid Intrinsic Disorder Analysis Online (RIDAO), a disorder prediction software which integrates multiple intrinsic disorder prediction algorithms (Dayhoff & Uversky, 2023). RIDAO analysis revealed a heterogeneous disorder profile, with alternating predicted ordered and disordered regions (**Supplementary Fig. 1B**). Notably, AP2-L contained a central region (residues 550-940) with high predicted disorder, with multiple algorithms assigning scores above the disorder threshold of 0.5. This region was enriched in low-complexity, polar and charged residues, including asparagine, serine, glycine, aspartate, and glutamate, consistent with the sequence characteristics of intrinsically disordered regions (IDRs). We next examined a region (residues 550–940) corresponding to a major RIDAO-predicted disordered region, using PSIPRED (McGuffin et al., 2000) to assess secondary-structure propensity. Although this region was predominantly predicted to adopt a coil conformation, discrete α-helical and β-strand segments were also identified. Thus, despite its intrinsic disorder propensity, this portion of AP2-L contains localized sequence elements with the potential to adopt secondary structure. Because IDRs can mediate protein interactions through short molecular recognition features (MoRFs) that undergo disorder-to-order transitions upon binding, we examined AP2-L sequence using MoRFpred, a sequence-based predictor of putative disorder-to-order transitioning binding regions (Disfani et al., 2012). MoRFpred did not identify high-confidence MoRFs within the major predicted IDR, suggesting that this region does not contain prominent canonical short binding elements predicted to undergo disorder-to-order transitions.

### AP2-L is likely essential for sporozoite infection *in vivo*

We generated marker-free parasites missing the AP2-L open reading frame (AP2-L KO) (**Supplementary Fig. 2A-B**). The growth rate of AP2-L KO asexual blood stages was determined by daily quantification of parasitemia in mice. Results of these experiments (**Supplementary Fig. 2C**) confirmed previous reports that AP2-L is dispensable for the intraerythrocytic cycle (Iwanaga et al., 2012) (Modrzynska et al., 2017). AP2-L KO parasites were transmitted to mosquitoes, and the number of sporozoites in mosquito SGs was determined in bulk. These data demonstrated that AP2-L KO parasites are infectious to mosquitoes and develop into SG sporozoites (**Supplementary Fig. 2D**).

Next, we characterized the ability of AP2-L KO parasites to undergo pre-erythrocytic development in the liver. Mice were intravenously infected with KO or Control sporozoites and monitored for the appearance of parasites in the bloodstream (**Table 1**). None of the KO-infected mice developed detectable blood stage parasitemia for 21 days whereas Control mice were positive in 6 days (**Table 1**). A second cohort of mice was infected through mosquito bite to mimic the natural route of infection (**Supplementary Fig. 2E**). In this case as well, KO-infected mice did not develop blood stage parasites while all Control-infected mice became patent. These data confirm previous results obtained using pooled mutants (Modrzynska et al., 2017). Since AP2-L is dispensable for the erythrocytic cycle (**Supplementary Fig. 2C**), the lack of patency by AP2-L KO sporozoites suggests that AP2-L function is critical for the parasite to infect, develop and/or exit infected hepatocytes *in vivo*.

**Table 1.** AP2L-KO sporozoites do not infect mice. Prepatent period was determined in mice inoculated with sporozoites.

| Parasite | Number of injected (IV) sporozoites | Number of mice injected | Number of patent mice at 3 weeks p.i. | Prepatent period ± SD (days) |
| --- | --- | --- | --- | --- |
| Control | 5000 | 8 | 8 | 5.9 ± 0.8 |
| KO | 5000 | 8 | 0 | No infection |

### AP2-L regulates the expression of invasion determinants

To determine the molecular function of AP2-L, we used bulk RNAseq to compare the transcriptomes of Control and AP2-L KO sporozoites. Principal component analysis revealed that the transcriptomes clustered according to parasite genotype (**Supplementary Fig. 3A**). Loss of AP2-L resulted in decreased abundance of 251 transcripts and increased abundance of 363 transcripts (|log_2_fold change| greater than 1, false discovery rate of 5%). The most robust and statistically significant transcriptional shifts occurred among transcripts that were less abundant in KO sporozoites, suggesting that AP2-L may act primarily to promote the production or stabilization of transcripts rather than to repress transcription or increase transcript degradation (**Fig. 1A**). A Gene Ontology (GO) analysis of transcripts that were significantly reduced in AP2-L KO sporozoites showed enrichment for biological process terms associated with movement in host environment (4-fold, 21 genes), entry into host (4-fold, 15 genes) and interaction with the host (3.5-fold, 21 genes) (**Fig. 1B**). Consistent with these functional categories, cellular component (CC) analysis revealed strong enrichment for proteins localized to the cell surface (24 genes), micronemes (15 genes) and apical part of the cell (23 genes) (**Supplementary Table 1**). When a more stringent cutoff of |log2 fold change| greater than 2 was applied, 48 transcripts were found to be less abundant in AP2-L KO sporozoites. GO analysis of this smaller subset identified enrichment of processes associated with cell motility, locomotion, and carbohydrate and organic-acid metabolism (**Supplementary Table 1)**.

**Figure 1.**
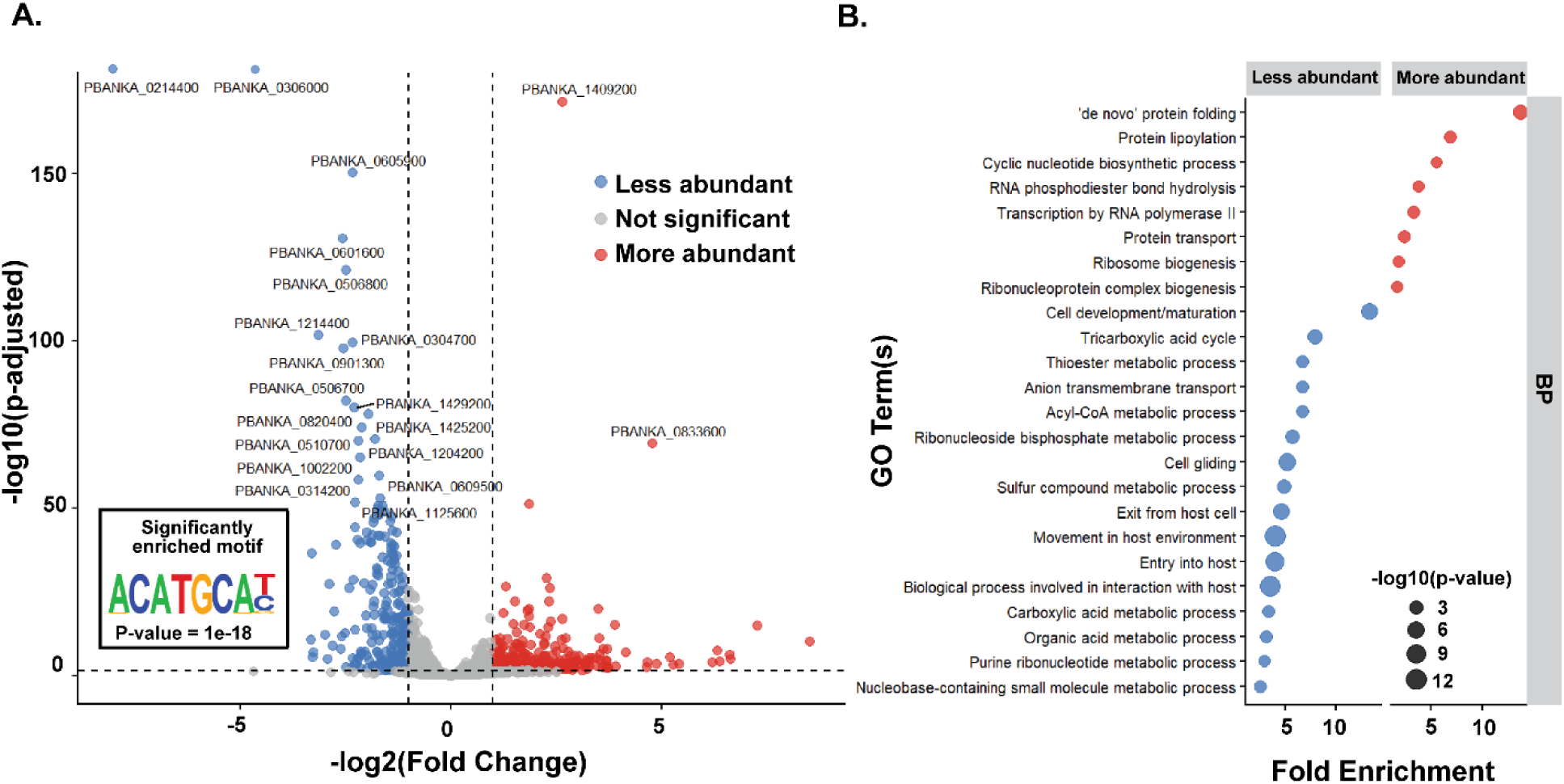
AP2-L functions as a transcriptional activator of genes involved in sporozoite-host interactions and liver-stage development. **(A)** Volcano plot showing differential gene expression in salivary gland sporozoites from AP2-L KO relative to Control. Each point represents a single gene. Genes with an absolute log_2_ fold change greater than 1 and an adjusted P-value less than 0.05 were considered differentially expressed. The inset shows the sequence logo and enrichment P-value of an enriched motif upstream the transcription start sites of significantly downregulated genes **(B)** Gene Ontology (GO) enrichment analysis of biological process terms among the significantly more and less abundant genes in AP2-L KO sporozoites.

To identify potential direct molecular targets of AP2-L, we conducted an *in-silico* motif enrichment search of the 5’ untranslated regions (UTR) of transcripts whose abundance was affected in the KO. This analysis revealed a two-fold (56% vs 29%) enrichment of a CATGCA-centered motif -1000 to +300 bp relative to the transcription start site among transcripts that were significantly less abundant in KO compared to non-differentially expressed genes (**Fig. 1A, Supplementary Table 1**). We reasoned that these genes represent candidate AP2-L-regulated targets. GO analysis of the motif-containing subset of genes identified biological process terms associated with interaction with the host (4-fold), cell motility (4.21-fold), entry into the host (4.06-fold), and exit from the host cells (5.75-fold). Cellular component (CC) analysis of the same subset revealed enrichment for micronemes (9.02-fold), the cell surface (8.39-fold), and the apical part of the cell (3.28-fold), highlighting the representation of genes encoding proteins associated with the host-parasite interface (**Supplementary Fig. 3B**). Motif-containing genes included ones whose product are required for sporozoite gliding, cell traversal and/or hepatocyte infection, including Claudin-like apicomplexan microneme protein (CLAMP) (Loubens et al., 2023), CLAMP-linked invasion protein (CLIP) (Unhale et al., 2026), CSP (Tewari et al., 2002), Sporozoite surface protein essential for liver-stage development (SPELD) (Al-Nihmi et al., 2017), Thrombospondin-related sporozoite protein (Labaied et al., 2007), Perforin-like protein 1 (PLP1) (Tomoko Ishino et al., 2005) and LIMP (Santos et al., 2017). In addition, the CATGCA-centered motif was present in genes encoding proteins with established or proposed roles in sporozoite-host interactions, such as Sporozoite invasion-associated protein 1 (SIAP1) (Engelmann et al., 2009), TRAP-like protein (TLP) (Moreira et al., 2008), P52 (T. Ishino et al., 2005), Membrane associated erythrocyte binding-like protein (MAEBL) (Yang et al., 2017), merozoite surface protein 1 (Combe et al., 2009) and cysteine repeat modular protein 4 (Douradinha et al., 2011) (**Supplementary Table 1**). Several members of the 6-cysteine rich family, including P52, P36, and P38, were less abundant in AP2-L KO sporozoites and contained the motif (Fernandes et al., 2023) (van Schaijk et al., 2008). Together, these data point towards AP2-L-regulated processes playing a major role in parasite invasion of and development within the host liver. However, direct binding of AP2-L to the upstream regions of the candidate target genes remains to be tested.

### AP2-L is required for the formation of a robust replicative vacuole

To better understand the failure of AP2-L KO parasites to undergo intrahepatocyte development, we first examined host-cell entry by AP2-L KO sporozoites by quantifying the fraction of sporozoites that are intracellular at 2 h post-inoculation (p.i.) of HepG2 cells. A similar fraction of AP2-L KO and Control sporozoites became internalized when added to HepG2 cells, indicating that AP2-L related processes do not affect the parasite’s entry into hepatocytes **(Fig. 2A, Supplementary Fig. 4A)**. Sporozoites exhibit three forms of cell entry. First, forming a replication-competent vacuole within which they undergo morphological transformation into an EEF (Risco-Castillo et al., 2015) (Mota et al., 2001). Second, forming a transient, non-replicative vacuole during their migration through cells (Risco-Castillo et al., 2015). Third, entering without forming a vacuole leading to damage to the host cell plasma membrane (Mota et al., 2001). To differentiate the fraction of AP2-L KO sporozoites that were in replicative-competent vacuoles, we visualized sporozoites enveloped in the vacuole membrane protein UIS4 at 6 h p.i. At this time point, almost 100% of Control sporozoites were enveloped in strong, continuous UIS4 expression (category 1 and 2, **Fig. 2B, Supplementary Fig 4B**) signifying a robust PVM. The percentage of AP2-L KO sporozoites that were surrounded by a UIS4-positive vacuole was 70% (category 1 and 2, **Fig. 2B, Supplementary Fig 4B**). The remaining AP2-L KO parasites displayed patchy UIS4 expression (category 3, **Fig. 2B, Supplementary Fig 4B**). The membranes of these parasites appeared to be blebbing and their cytoplasm appeared fragmented or vesiculated. We interpret these observations to imply that these sporozoites were being degraded. We quantified the amount of UIS4 in the parasite using automated image analysis of sporozoites at 6 h p.i. and Western blotting of SG sporozoites (**Supplementary Fig 4D)**. UIS4 abundance is decreased in AP2-L KO parasites, suggesting a deficiency in a key protein component of the PVM. Together, these data suggest that AP2-L is critical for the optimal formation of a robust replicative vacuole.

**Figure 2.**
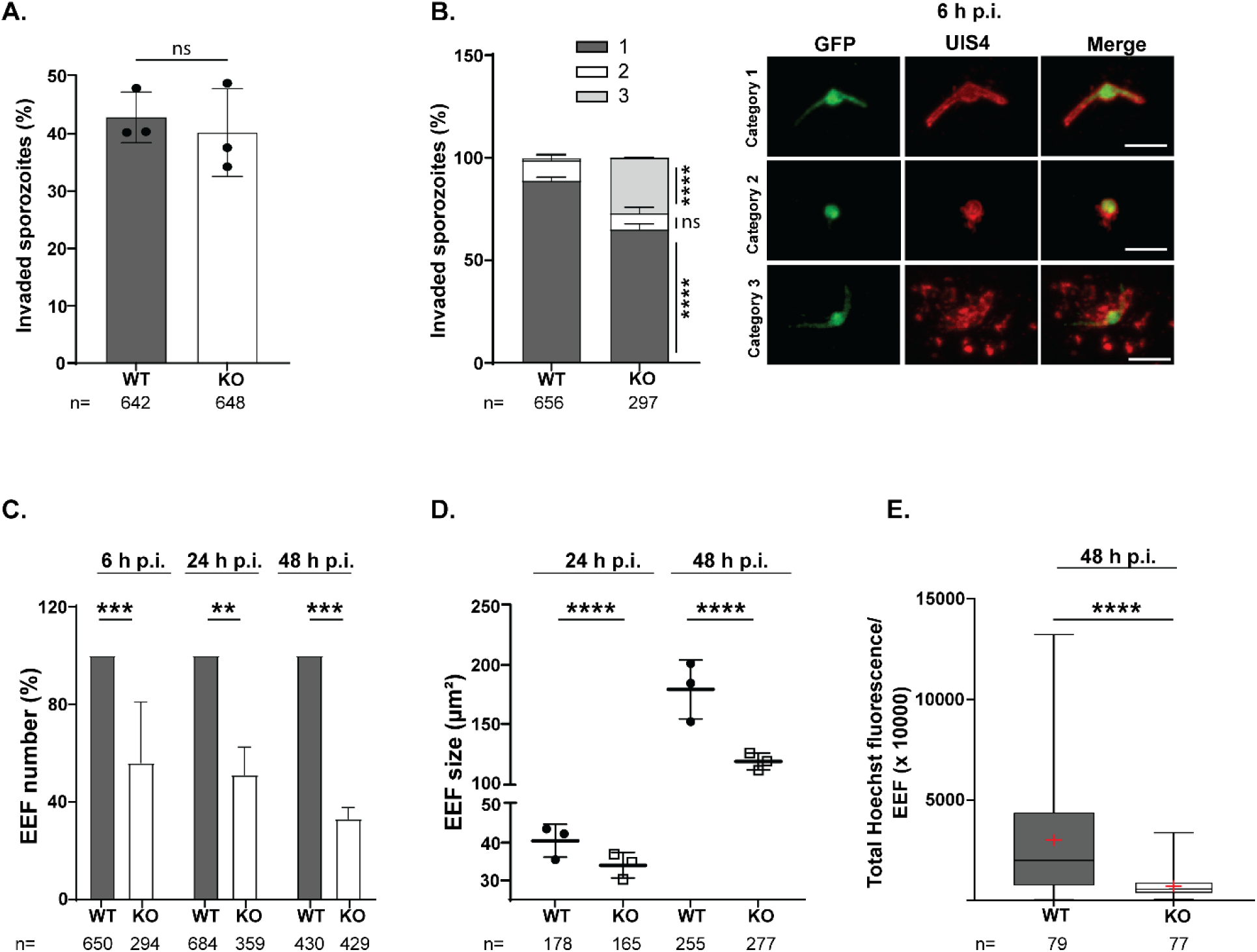
AP2-L is dispensable for hepatocyte invasion but is required for sporozoite transformation and liver-stage growth. **(A)** Invasion of HepG2 cells by Control and AP2-L KO sporozoites was quantified using double differential immunofluorescence staining. Invasion efficiency was calculated as the percentage of intracellular sporozoites relative to the total number of sporozoites. **(B)** Sporozoite transformation into EEFs was quantified at 6 h p.i. using GFP to visualize the parasite cytoplasm (green) and UIS4 to visualize the PVM (red). Parasites were assigned to one of three categories based on parasite morphology and UIS4 distribution. Representative images of the three categories are shown. Scale bars, 3 µm. **(C)** The number of EEFs was quantified at 6, 24, and 48 h p.i. **(D)** The size of EEFs was quantified at 24 and 48 h p.i. **(E)** DNA replication was assessed at 48 h p.i. by quantifying Hoechst fluorescence intensity within each GFP-defined EEF. Data represent the mean ± SD from three or more independent experiments. Statistical significance was determined using an unpaired two-tailed Student’s t-test.

### AP2-L is required for growth of EEFs

Next, we examined the effect of AP2-L loss on EEF formation and growth. We quantified the number and size of EEFs present in culture at 6-48 h p.i. (**Fig. 2C-D, Supplementary Fig. 4E-F)**. There were approximately 50% fewer AP2-L KO EEFs at 24 h p.i., and they were significantly smaller in size (14%) compared to Control, demonstrating a defect in sporozoite transformation. This growth defect was more pronounced at 48 h p.i., with KO EEFs being 46% smaller than Control, suggesting a defect in trophozoite growth (**Fig. 2C-D, Supplementary Fig. 4E-F)**. To determine if there was an effect on transitioning from the trophozoite to the hepatic schizont stage, we quantified relative DNA content by measuring total Hoechst fluorescence within individual GFP-defined EEFs. AP2-L KO EEFs exhibited significantly lower relative DNA content than Control parasites (**Fig. 2E**). These data suggest that while AP2-L-dependent processes are not required for the initiation of nuclear division, they play a critical role in the continuation of the parasite’s hepatic schizogony.

We reasoned that the decrease in size of KO EEFs at 24-48 h p.i. could result from an impairment of the PVM, as was suggested by the aberrant UIS4 localization in KO parasites at 6 h p.i. (**Fig. 2B, Supplementary Fig 4B).** We examined the localization of UIS4 and another PVM protein, EXP1 (Spielmann et al., 2012) in developing EEFs, with GFP used to delineate the parasite cytoplasm. At 24 h p.i., nearly all control EEFs displayed contiguous UIS4 localization throughout the PVM and in the TVN extensions projecting into the host-cell cytoplasm **(Fig. 3A, Supplementary Fig. 5A-B)**. EXP1 was detected within the cytoplasm at 24 h p.i. and prominently localized to the PVM by 48 h p.i. marking these EEFs as mid-stage **(Fig. 3B, Supplementary Fig. 5C).** In contrast, at 24 h p.i., 60% of KO EEFs displayed patchy UIS4 localization in the PVM and at 48 h p.i., only 36% of KO EEFs had any detectable UIS4 in the PVM **(Fig. 3A, Supplementary Fig. 5A-B)**. In the remaining KO EEFs, UIS4 was detected only in the cytoplasm. The KO PVM appeared hypervessiculated and discontinuous. There were almost no TVN structures in KO-infected cells but UIS4-positive puncta were present around the EEF body in 61% of AP2-L KO parasites. These could represent remnants of the PVM or TVN extensions formed by the KO EEF (**Fig. 3A, Supplementary Fig. 5B)**. In addition, KO EEFs did not display EXP1 expression at either time-point **(Fig. 3B, Supplementary Fig. 5C)**, consistent with a stall at the early-to-mid hepatic schizont stage and the reduction in relative DNA content at 48 h p.i. (**Fig. 2E)**.

**Figure 3.**
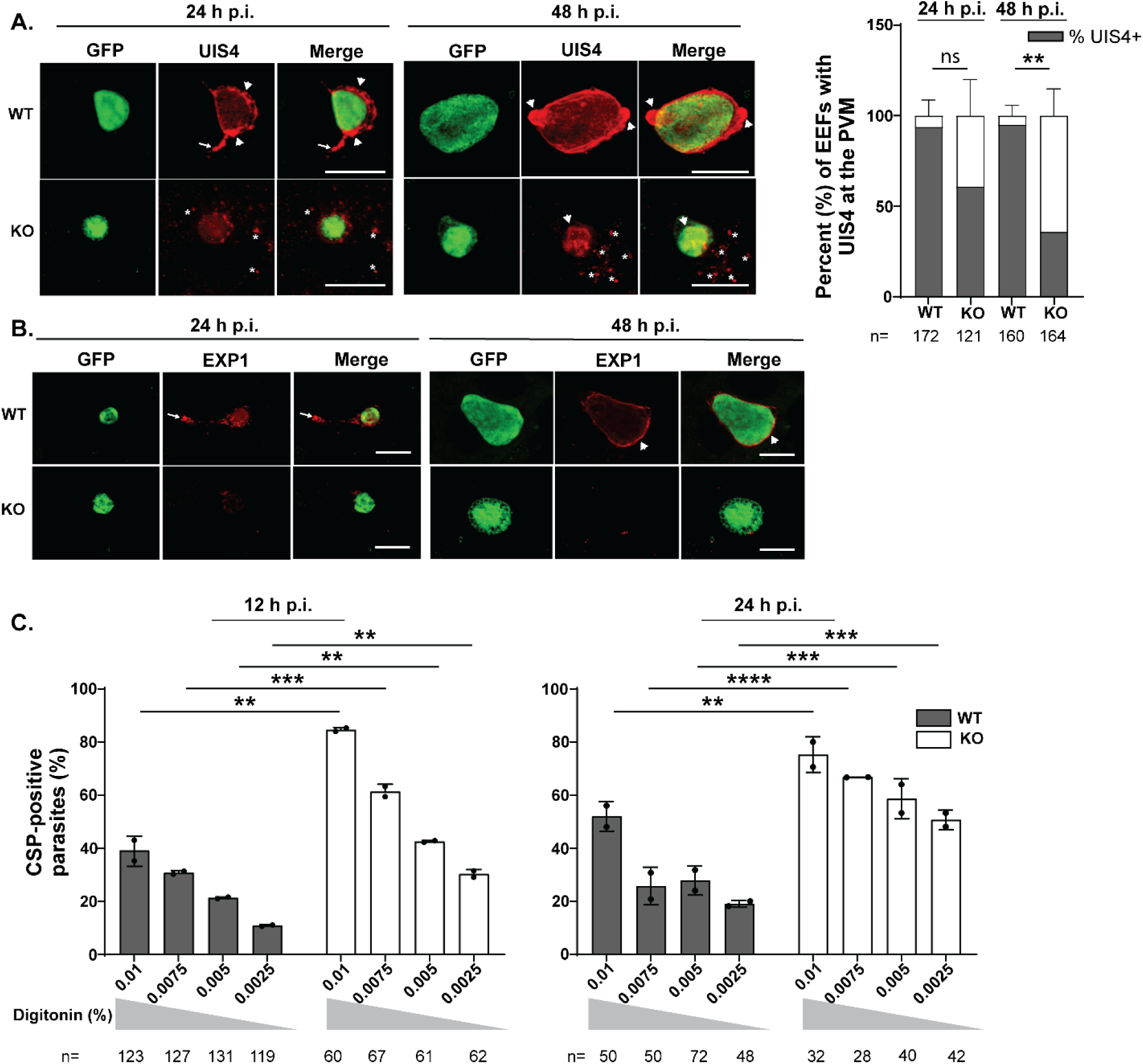
Loss of AP2-L disrupts localization of PVM proteins and compromises PVM integrity. **(A)** Representative immunofluorescence images of HepG2 cells infected with Control or AP2-L KO sporozoites with the parasite cytoplasm visualized using GFP and the PVM visualized using UIS4. Arrowheads indicate the PVM surrounding the parasite, arrows indicate PVM extensions and asterisks indicate UIS4-positive puncta. Scale bars, 10 µm. The graph shows the percentage of EEFs displaying detectable UIS4 localization at the PVM. Statistical significance was determined using an unpaired two-tailed Student’s t-test (ns, P> 0.05; **P ≤ 0.01). **(B**) Representative immunofluorescence images of EEFs with the parasite cytoplasm visualized using GFP and the PVM visualized using EXP1. Arrowheads indicate the PVM surrounding the parasite, and arrows indicate PVM extensions. Scale bars, 10 µm. **(C)** Digitonin permeabilization assay was used to assess PVM porosity. The percentage of CSP-positive parasites was quantified at different digitonin concentrations. Data represents the mean ± SD from 2 or more independent experiments. Statistical significance was determined using 2-way ANOVA (**P ≤ 0.01; ***P ≤ 0.001; ***P ≤ 0.0001).

UIS4 mislocalization in KO EEFs suggests that AP2-L-dependent processes are required for protein trafficking to the PVM and/or maintenance of the PVM. To further examine the integrity of the PVM in KO parasites, we developed an assay for PVM porosity (**Fig. 3C, Supplementary Fig. 5D**). Infected HepG2 cells were treated with increasing concentrations of the non-ionic detergent, digitonin, to differentially permeabilize the host-cell plasma membrane and the PVM (Marques-da-Silva et al., 2025). A permeabilized PVM allows access to antibodies that recognize the parasite plasma membrane protein, CSP, while an intact PVM excludes them (**Supplementary Fig. 5D)**. At the highest digitonin concentrations, nearly all control and KO EEFs displayed anti-CSP staining, demonstrating permeabilization of their PVM. In contrast, at lower digitonin concentrations, a significantly higher proportion of KO EEFs were CSP-positive compared to control (**Fig. 3C**). The difference was seen as early as 12 h p.i. (3-fold higher anti-CSP reactivity in KO versus control at the lowest digitonin concentration), demonstrating that the KO PVM is compromised from the start. These data are consistent with loss of AP2-L compromising the PVM, making it more sensitive to low levels of detergent. The loss of membrane integrity could arise from altered protein and/or fatty acid composition of the PVM.

### AP2-L-dependent processes are required for evasion of host autophagy

The progressive reduction in KO EEF numbers despite their ability to enter host cells, together with the discontinuous localization of UIS4 in the PVM and vesiculated appearance of the cytoplasm of the EEF, suggested that KO parasites were undergoing clearance by the host cell. We therefore examined the effects of AP2-L-dependent processes on the protective function of the PVM by characterizing LC3 accumulation around the KO EEF. Hepatocytes mount a cytosolic response to intracellular sporozoites through rapid recruitment of the autophagic marker LC3 to the PVM (Prado et al., 2015). Successful parasite development requires redistribution of LC3 away from the parasite body into the TVN extensions and its eventual shedding into the host cytoplasm. We quantified LC3 fluorescence around the PVM at 12 and 24 h p.i. using automated image analysis (**Supplementary Fig. 6A**). At 12 h p.i., there was no significant difference in the amount of LC3 associated with KO and control parasites normalized for either parasite-body size or total cellular LC3 (**Fig. 4A; Supplementary Fig. 6B)**. However, at 24 h p.i., KO EEFs accumulated significantly more LC3 compared to control **(Fig. 4A; Supplementary Fig. 6C).** Since lipidated LC3 attachment to the PVM is essential for the formation of the autophagosome, healthy parasites shed LC3 to avoid elimination. The persistent LC3 association with KO parasites at 24 h p.i. indicates enhanced autophagy-related targeting and suggests that these EEFs are unable to prevent clearance by the host cell.

**Figure 4.**
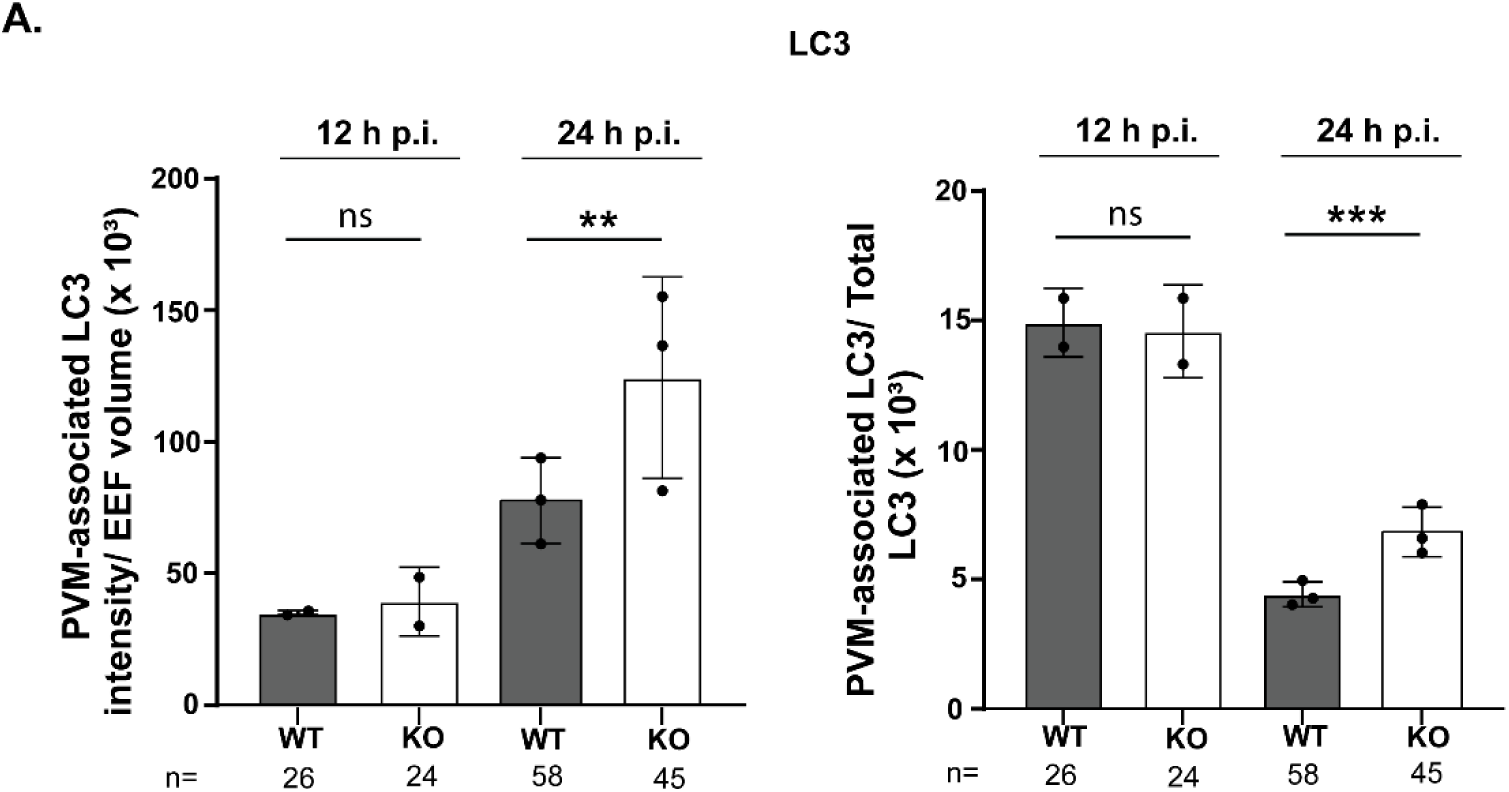
Loss of AP2-L increases LC3 accumulation around the PVM. LC3 fluorescence within a 2 µm region surrounding the PVM was quantified. The PVM-associated LC3 signal was normalized to EEF volume (left) and to total cellular LC3 (right). Data represent the mean ± SD from 3 or more independent experiments. Statistical significance was determined using an unpaired two-tailed Student’s t-test (ns, P > 0.05; **P ≤ 0.01; ***P ≤ 0.001).

### AP2-L-dependent processes affect the acquisition of host vesicular traffic

Another key function of the PVM is nutrient uptake from the host. We reasoned that the smaller size of KO EEFs could result from nutrient deficiencies caused by its distorted PVM. ApoH, a host apolipoprotein required for efficient liver-stage growth, is thought to be recruited through interactions with cytoplasmic domain of EXP1 (Sa et al., 2017). We quantified ApoH accumulation around individual EEFs at 24 h p.i. using automated image analysis and normalized the signal to EEF size (**Supplementary Fig. 6D**). Despite the reduction in EXP1 localization in KO parasites, there was no significant difference in ApoH accumulation around control and KO EEFs **(Supplementary Fig. 6E)** suggesting that EXP1-independent pathways may be compensating for the loss of EXP1-based uptake of ApoH in KO EEFs.

Another mechanism employed by the parasite to acquire host-derived nutrients is through exploitation of the host vesicular trafficking machinery (Atchou et al., 2026) (Vijayan et al., 2022) (Raphemot et al., 2019). The parasite repositions the host’s microtubule organizing center (MTOC) from its typical juxtanuclear position to the PVM, enabling redirection of the vesicular traffic from the host Golgi towards itself (Vijayan et al., 2022). We examined this process in AP2-L KO parasites by first assessing their position relative to the host nucleus from 6 to 48 h p.i. To account for the smaller size of AP2-L KO EEFs, the PVM-to-host nucleus distance was normalized to EEF volume to account for differences in parasite size. Control EEFs moved progressively closer to the host nucleus as they matured (**Fig. 5A**). At 6 h p.i., KO and control parasites were positioned similarly relative to the host nucleus, with mean normalized distances of 0.12 µm and 0.10 µm, respectively (**Fig. 5A, Supplementary Fig. 7A**). However, beginning 12 h p.i. AP2-L KO EEFs exhibited a significantly greater normalized distance from the host nucleus (0.09 µm) than control EEFs (0.01 µm) and this difference persisted at later time points (0.02 µm vs 0.008 µm at 24 h p.i. and 0.003 µm vs 0.0004 µm at 48 h p.i., KO vs Control, respectively; **Fig. 5A, Supplementary Fig. 7A**).

**Figure 5.**
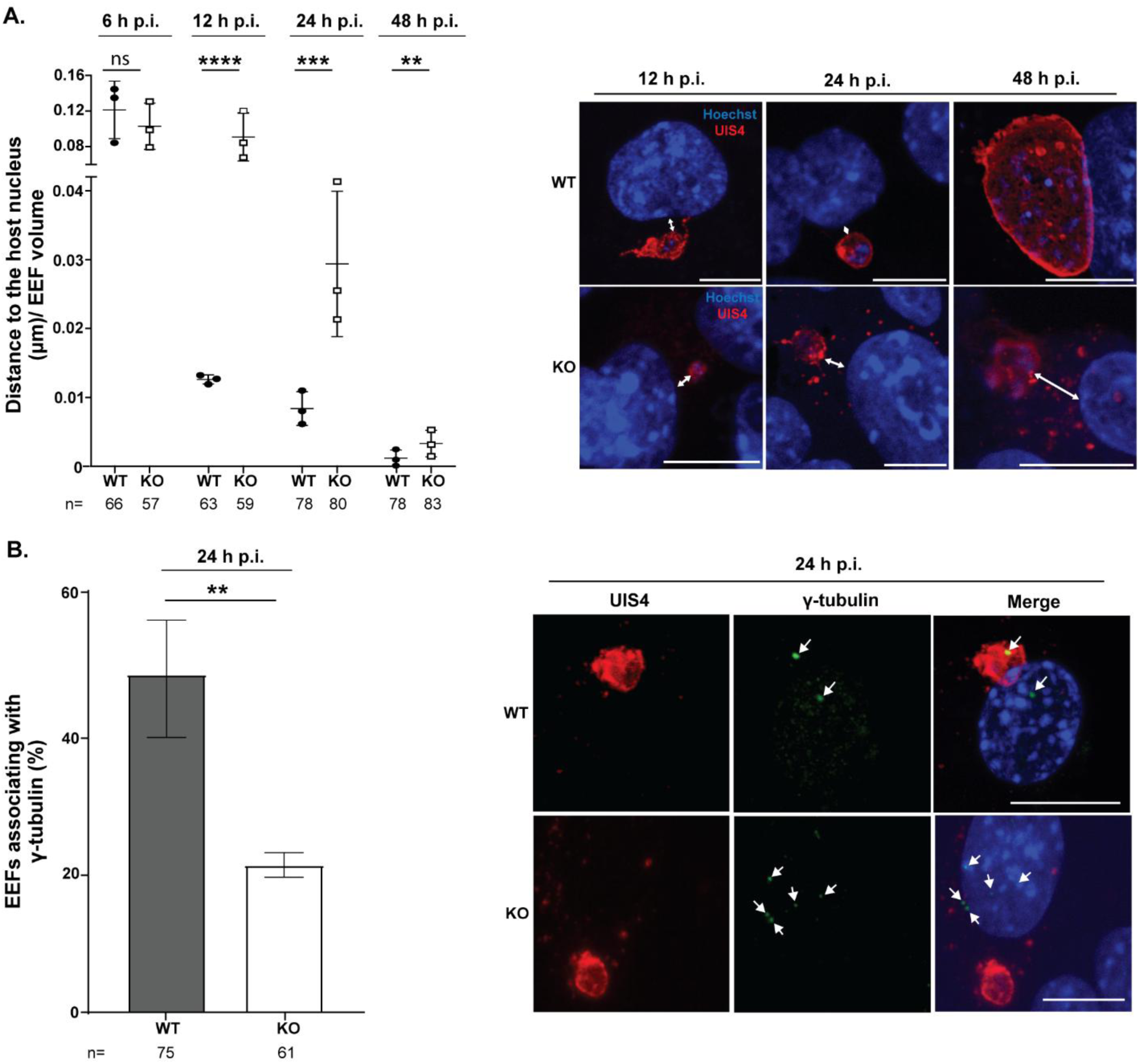
Loss of AP2-L alters EEF perinuclear positioning and reduces association with host γ-tubulin foci. **(A)** Infected HepG2 cells were stained for GFP, UIS4, host, and parasite nuclei. Maximum-intensity projections were used to measure the distance between UIS4-positive PVM and the periphery of the host-cell nucleus. This distance was normalized to EEF volume. Representative immunofluorescence images show parasite relative to the host nucleus at 12, 24, and 48 h p.i. **(B)** The percentage of EEFs associated with host γ-tubulin foci at 24 h p.i. Representative immunofluorescence images show UIS4-positive EEFs (red), γ-tubulin (green, arrows), and Hoechst-labeled nuclei. Data represent the mean ± SD from 3 independent experiments. Statistical significance was determined using an unpaired two-tailed Student’s t-test (ns, P > 0.05; **P ≤ 0.01; ***P ≤ 0.001; ****P ≤ 0.0001). Scale bars, 10 µm.

We reasoned that a host nucleus-adjacent position could assist the parasite in reorienting the MTOC towards the PVM. We examined the position of the MTOC relative to the PVM by visualizing γ-tubulin foci and UIS4 in infected cells. At 24 h p.i., the MTOC was found associated with the PVM of 48.03 % of control EEFs and 21.52 % of AP2-L KO EEFs (**Fig. 5B, Supplementary Fig. 7B)**. Next, we examined if the inefficient capture of the MTOC by KO EEFs affects the parasite’s ability to re-direct the host vesicular traffic. We visualized the contact between the parasite and the host Golgi using the cis-Golgi marker, GM130 and the trans-Golgi marker, Golgin97 (**Fig. 6A-B, Supplementary Fig. 7C-D)**. Contact between the host Golgi and the parasite was detected in approximately 80% of control and 40% of KO EEFs. Control parasites contacted the host Golgi through both the PVM and TVN extensions (**Fig. 6A-B, Supplementary Fig. 7C-D)**, while contact by KO EEFs, when detectable, occurred through the PVM. Vesicles transiting through the endomembrane system can be visualized using COPB2 (Raphemot et al., 2019). COPB2 staining showed significantly higher recruitment of vesicles to the PVM of control EEFs compared to KO EEFs (81% vs 42%) (**Fig. 6C, Supplementary Fig. 7E**). Together, these results suggest that AP2-L-dependent processes are required for MTOC capture and subsequent hijacking of vesicular traffic emanating from the host Golgi. Impairment of this process would limit acquisition of host lipids and other nutrients required for EEF membrane formation and growth, thereby limiting EEF size.

**Figure 6.**
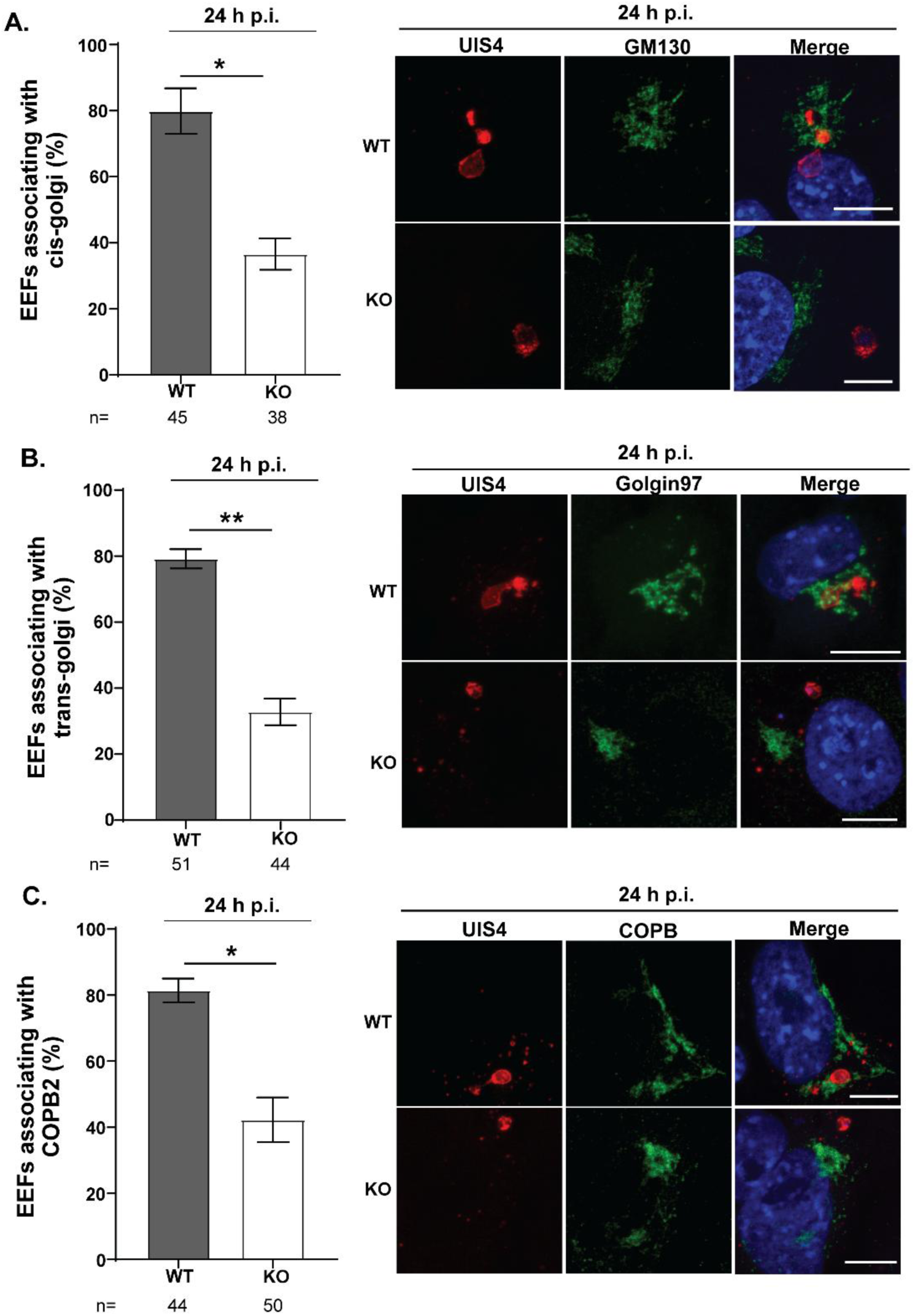
Loss of AP2-L alters interactions with host vesicular pathways. **(A)** Percentage of EEFs associated with the cis-Golgi marker, GM130. Representative immunofluorescence images show UIS4, GM130 and the merged channels with Hoechst. **(B)** Percentage of EEFs associated with the trans-Golgi marker, Golgin-97. Representative immunofluorescence images show UIS4, Golgin-97 and the merged channels with Hoechst. **(C)** Percentage of EEFs associated with COPB2-positive structures. Representative immunofluorescence images show UIS4, COPB2, and the merged channels with Hoechst. Data represents the mean ± SD from 3 independent experiments. Statistical significance was determined using an unpaired two-tailed Student’s t-test (*P ≤ 0.05; **P ≤ 0.01). Scale bars, 10 µm.

### AP2-L KO EEFs have an intrinsic developmental defect

EEF development in the liver is asynchronous, and parasites mature at different rates. It is possible that some AP2-L KO EEFs are capable of completing development albeit slowly but are eliminated by the host’s innate responses. Interferon (IFN) signaling is upregulated early in liver-stage development and acts to control the development of parasites in the liver (Liehl et al., 2015) (Miller et al., 2014). We tested the ability of AP2-L KO sporozoites to infect mice lacking the type I and III IFN signaling pathways. We infected Control and mutant mice with KO sporozoites and monitored the appearance of blood stage parasites for 21 days (**Supplementary Table 2**). None of the mice developed infection in this period. These data suggest that the infection defect resulting from the loss of AP2-L is parasite-intrinsic and not due to more efficient clearance of slow-growing EEFs by innate immunity.

## Discussion

We corroborate previous findings that *P. berghei* AP2-L is dispensable for asexual stage growth, sporozoite formation, and hepatocyte invasion but essential for LS growth. In addition, as novel observations, we find that AP2-L-dependent processes are required for PVM formation/maintenance and nuclear replication in the hepatic trophozoite. Our data suggest a model in which AP2-L directly or indirectly promotes the abundance of transcripts encoding several micronemal-resident proteins involved in PVM formation/maintenance (p36, p52, CLAMP, CLIP, SPATR, TRAP, CeLTOS and PLP-1) contribute to the formation of a robust, intact and continuous PVM. The PVM dictates interactions between the parasite and host cell, supporting parasite growth by resisting clearance by autophagy-like pathways, acquiring nutrients, and expelling waste metabolites. We hypothesize that the PVM has an additional, as-yet unrecognized function in ensuring parasite proximity to the host nucleus, which may assist in MTOC capture and subsequent interaction with the host’s vesicular traffic. We were surprised to see normal invasion by AP2-L KO sporozoites despite reduced abundance of TRAP, CS and TLP (Sultan et al., 1997) (Tewari et al., 2002) (Moreira et al., 2008). One explanation is that while our studies in tissue-culture cells did not reveal a defect in sporozoite invasion, it is possible that *in vivo* studies would detect changes in the KO sporozoite’s ability to exit the skin or invade hepatocytes.

The impairment in the parasite’s PVM as a result of AP2-L loss was suggested by aberrant UIS4 localization, its jagged appearance, the absence of a TVN in AP2-L KO-infected cells, and the increased sensitivity of the KO PVM to digitonin. The loss of PVM integrity in the AP2-L KO could result, at least in part, from defective rhoptry discharge since CLAMP, SPATR and CLIP transcripts were less abundant in KO sporozoites. These micronemal proteins are required for rhoptry discharge in *Toxoplasma gondii* (Valleau et al., 2023), and their loss in *P. berghei* sporozoites abrogates invasion (Loubens et al., 2023). Several rhoptry proteins are part of the tight junction and become incorporated into the PVM (Beckers et al., 1994; Ben Chaabene et al., 2021; Ghosh et al., 2017). Thus, a collective impairment in the release of rhoptry proteins, as a cumulative consequence of decreased abundance of CLAMP, SPATR and CLIP, could reasonably be expected to impact PVM integrity. Other micronemal proteins whose transcripts were less abundant in AP2-L KO sporozoites include p36 and p52, both members of the 6-cysteine protein family that have been implicated in PVM formation. Sporozoites lacking p36 and p52 are able to enter hepatocytes but do not form a PVM (Arredondo et al., 2018; Ploemen et al., 2012). Additional members of the 6-cysteine family with less abundant transcripts in AP2-L KO sporozoites are p230 and p38. While mutant sporozoites with deletions in either gene (van Dijk et al., 2010) (Lin et al., 2011) (Fernandes et al., 2023) have no reported defects in the differentiation of sporozoites into liver stages, a decrease in their expression along with a decrease in p36 and p52 could have the cumulative effect of compromising PVM integrity. Future work will focus on assessing the effect on 6-cysteine proteins and their associated complexes in KO EEFs to test this model.

Even when AP2-L KO parasites do form a PV and transform into EEFs, their growth is slower than Control, likely due to nutritional deficiencies that can limit the formation of the TVN. Nutrients required by the growing EEF are lipoic acid (Deschermeier et al., 2012), phospholipids, such as phosphatidyl ethanolamine and phospatidylcholine (Mikolajczak et al., 2007) (Sharma et al., 2008) (Itoe et al., 2014) and metal ions like zinc and iron (reviewed in (Nyboer et al., 2018), (Sahu et al., 2014). The normal accumulation of ApoH in AP2-L KO parasites despite the lack of EXP1 suggests that there are multiple routes for the EEF to acquire host lipids. One such route may be directly from the host ER via membrane contact sites between it and the PVM (Liebeck et al., 2026) (Romano et al., 2025).

Another implication of our work is that the extended amino-terminal region of AP2-L contributes to protein function. Neither of the two AP2 domains of *P. falciparum* AP2-L displayed strong DNA-binding affinities *in vitro* (Campbell et al., 2010), although a tandem-domain construct was associated with an AATTTCC recognition motif. Computational searches for enriched motifs upstream of differentially regulated genes in EEFs lacking the AP2 C-terminal DNA-binding region did not reach conclusive results (Iwanaga et al., 2012). More broadly, not all ApiAP2 proteins appear to function solely through sequence-specific DNA binding. There is a precedent for functionally important protein-protein interactions mediated by regions outside the AP2 domain. For example, *P. falciparum* AP2-HC associates with heterochromatin independently of its AP2 domain via interaction with heterochromatin protein 1 (*Pf*HP1) (Carrington et al., 2021). ApiAp2 proteins have also been identified in complexes with chromatin-modifying and transcriptional regulatory proteins in *Toxoplasma gondii* (Wang et al., 2014).

One possible function of the amino-terminal region of *P. berghei* AP2-L may be refining the promoter binding specificity of the AP2 domains. In higher eukaryotes, IDRs of transcription factors have been reported to act as scaffolds for the general transcriptional machinery, acting to concentrate the transcriptional machinery in phase-separated condensates and/or affecting promoter selectivity *in vivo* (Strugo et al., 2025) (Brodsky et al., 2021) (Brodsky et al., 2020) (Boija et al., 2018). Concordantly, the phenotypes associated with the complete ap2-l deletion described here are broader than those reported following disruption of only the C-terminal region (Iwanaga et al., 2012). The latter found *P. berghei* AP2-L was not essential for LS development since sporozoite-infected rodents eventually developed blood stage parasitemia. Disruption of the C-terminal region resulted in reduced EEF size but did not significantly affect EEF number or UIS4 localization, and expression analysis of selected genes in SG sporozoites did not reveal substantial differences. It is possible that a truncated protein with partial activity is made in these mutants.

Our *de novo* HOMER analysis of genes whose transcript were less abundant in KO sporozoites identified significant enrichment of a CATGCA-centered sequence. This sequence closely resembles the TGCATGCA motif recognized by AP2-SP in rodent malaria parasites (Yuda et al., 2010) and its *P. falciparum* ortholog, AP2-EXP (De Silva et al., 2008), with AP2-EXP binding to this motif further supported by ChIP-seq (Russell et al., 2022). This differs from the AATTTCC motif attributed to the recombinantly expressed tandem AP2 domains of *P. falciparum* AP2-L (Campbell et al., 2010). A targeted HOMER analysis found no significant enrichment of AATTTCC among genes whose transcripts were less abundant (33.1% of target vs 31.7% of background) in AP2-L KO sporozoites. Although the AP2 domains of *P. berghei* and *P. falciparum* AP2-L are conserved, they are not identical, and these sequence differences could contribute to species-specific differences in DNA-binding specificity. Discrepancies between *in vitro* binding preference of isolated AP2 domains and *in vivo* genomic localization of APIAP2 proteins has been reported previously. For example, the CACACA motif, preferentially bound by the recombinant *Pf*AP2-HC AP2 domain (Campbell et al., 2010), showed no positional association with *Pf*AP2-HC *in vivo* (Carrington et al., 2021). Nevertheless, the enrichment of the CATGCA motif in subset of genes whose transcripts were less abundant in AP2-L-KO sporozoites cannot presently be interpreted as evidence of AP2-L binding. Future work will focus on direct experimental validation using biochemical and genomic approaches such as ChIP-seq or CUT&RUN.

## Methods

### Ethics statement

All animal procedures in this project were reviewed and approved by the Institutional Animal Care and Use Committee (IACUC) of Rutgers, New Jersey Medical School. Experiments were conducted in accordance with the Animal Welfare Act, The Institute of Laboratory Animal Resources Guide for the Care and Use of Laboratory Animals and Public Health Service Policy.

### Experimental animals, parasite line and maintenance

Female Swiss Webster (S/W) mice (6-8 weeks old) were used in this study. *P. berghei* (strain ANKA) parasites constitutively expressing green fluorescent protein and luciferase (*Pb*-GFP-Luc) were used as the wild-type line. Female *Anopheles stephensi* mosquitoes were obtained from the Center for Global Infectious Disease Research at Seattle Children’s Hospital (Seattle, WA). Mosquitoes were maintained on filtered tap water containing 10% dextrose supplemented with 0.05% para-aminobenzoic acid (PABA) at 22-25°C and 75-80% relative humidity.

### Generation and validation of the *P. berghei* ap2-l knockout line

Marker-free parasites missing the AP2-L (PBANKA_0214400) open reading frame (AP2-L KO) were generated by inducing marker recombination in the RMgm line, RMgm-1733 (Modrzynska et al., 2017). Parasites were treated with 5-fluorocytosine (5-FC) to select parasites in which the marker cassette was recombined out. Individual parasite lines were obtained by limiting dilution.

### Mosquito infection and SG sporozoite isolation

Mosquitoes were fed on *P. berghei* infected S/W mice exhibiting high gametocytemia on the first two days and on uninfected mice on the third day. Mice were anesthetized by intraperitoneal injection of a cocktail containing 16 mg/ml ketamine and 1.2 mg/ml xylazine prepared in sterile 1x PBS. During the feeding period, mosquitoes were maintained with water containing 10% dextrose and 300µg/ml gentamicin solution. SG dissections were performed 21 or 22 days after the first blood meal.

### *In vivo* infections

Mice were inoculated intravenously with 5 x 10^3^ Control or AP2-L-KO sporozoites per mouse and monitored daily for 21 days beginning 24 h p.i. Blood stage parasitemia was assessed by detection of GFP-expressing parasites by flow cytometry. For flow cytometry analysis, a drop of tail blood was collected into RPMI 1640 supplemented with heparin. Samples were analyzed using a FACSymphony (BD Biosciences) flow cytometer. Data were analyzed using BD FACSDiva software. For bite transmission experiments, anesthetized mice were placed on top of mesh-covered mosquito cage containing approximately 50 infected mosquitoes for 5 min. Mice were monitored daily for 21 days by flow cytometry.

### HepG2 cell culture and sporozoite infection

Human hepatoma HepG2 cells were maintained in DMEM supplemented with 10% fetal bovine serum and 1x penicillin-streptomycin solution (cDMEM). Cells were cultured at 37°C in a humidified incubator containing 5% CO_2_. For infection experiments, HepG2 cells were seeded either in collagen (Gibco) solution coated 8-well Lab-Tek chamber slides or on sterile collagen coated glass coverslips placed in 12-well plates. Cells were seeded 48 h prior to infection and were at least 90% confluent on the day of infection. Freshly isolated Control or AP2-L KO SG sporozoites were diluted in cDMEM and added to each well (5 x 10^4^ per well). Medium was replaced with fresh cDMEM at 4 h p.i. Infected cultures were maintained until experimental time points. At each time point, cells were washed with PBS and fixed with 4% (w/v) paraformaldehyde (PFA) for 15 min at RT. Fixed samples were subsequently processed for immunofluorescence staining.

### Sporozoite invasion assay

Invasion assays were performed as previously described (Sinnis et al., 2013). Briefly, HepG2 cells (≥90% confluent) were inoculated with Control or AP2-L KO sporozoites and incubated for 2 h at 37°C in 5% CO_2_. Cells were fixed with 4% PFA in PBS for 20 min, blocked with 3% BSA in PBS for 30 min, and incubated with mouse anti-CSP antibody (clone 3D11; 1 µg/mL) for 1 h. After washing, cells were incubated with Alexa Fluor 594-conjugated anti-mouse secondary antibody for 1 h. Cells were then permeabilized with 100% cold methanol for 20 min, blocked again, and incubated with mouse anti-CSP antibody (clone 3D11; 1 µg/mL) followed by Alexa Fluor 488-conjugated anti-mouse secondary antibody. Nuclei were stained with Hoechst 33342 (1 µM), and coverslips were mounted with ProLong Diamond Antifade Mountant. Slides were stored at 4°C in the dark until imaging. Samples were imaged by fluorescence microscopy (Olympus BX61) using a 100x objective. At least 100 fields were examined per condition in at least three independent biological experiments. Intracellular sporozoites were determined by subtracting the number of Alexa Fluor 594-positive extracellular sporozoites from the total number of Alexa Fluor 488-positive sporozoites. Invasion efficiency was calculated as the percentage of total sporozoites that were intracellular.

### Indirect immunofluorescence assays

Infected cells were fixed with 4% PFA in 1x PBS for 15 min at RT. Cells were washed with PBS, permeabilized with 0.1% Triton X-100 (Sigma-Aldrich) for 30 min at RT and blocked with 3% BSA in PBS (blocking solution) for 1 h at RT. Cells were incubated with primary antibodies diluted in blocking solution overnight at 4°C. Information regarding the primary antibodies used, including target, host species, supplier, catalog number and dilution, is provided in **Supplementary Table 4**. Following primary antibody incubation, cells were washed three times with 1x PBS and incubated with AlexaFluor-conjugated secondary antibodies diluted in blocking solution for 3 h at RT in the dark. Cells were washed three times with PBS and nuclei were stained with Hoechst 33342 (1 µM) for 20 min at RT. After a final wash with PBS, a drop of ProLong Diamond Antifade Mountant was added to the air-dried slides before applying coverslips. Slides were stored at 4°C in the dark until imaging.

### Western blot

Control and AP2-L-KO sporozoites were collected in cold DMEM supplemented with cOmplete Mini EDTA-free protease inhibitor (Roche). Sporozoites were pelleted by centrifugation at 18,000xg for 10 min at 4°C. The pellets were resuspended in 2x Laemmli buffer (Thermo Fisher Scientific) supplemented with protease inhibitor. Samples were sonicated for two cycles using a 15s pulse followed by a 30s pause at 40% amplitude and were subsequently heated at 95°C for 10 min. Proteins were separated by SDS polyacrylamide gel electrophoresis (SDS-PAGE) using a precast 4-12% polyacrylamide gradient gel (ExpressPlus PAGE Gel; GenScript) and Tris-MOPS-SDS running buffer (GenScript). Before the transfer, the gel was equilibrated for 10 min at RT with gentle agitation in transfer buffer containing 25 mM Tris, 192 mM glycine, 20% methanol, and 0.1% SDS, pH 8.3. Proteins were transferred to an Immun-Blot polyvinylidene difluoride membrane (Bio-Rad), preactivated in methanol, at 90 V for 50 min at 4°C with continuous stirring. Following transfer, the membrane was rinsed with 1x PBS and blocked with 3% BSA in 1x PBS for 30 min at RT with gentle agitation. The membrane was incubated overnight at 4°C with primary antibodies diluted in blocking buffer. The membrane was washed three times with 1x PBS containing 0.1% Tween 20 (PBS-T) and incubated with the appropriate HRP-conjugated secondary antibody at a dilution of 1:20,000 for 1 h at RT. After three washes with PBS-T, chemiluminescent signal was developed by incubating the membrane with SuperSignal West Femto Maximum Sensitivity Substrate (Thermo Fisher Scientific) for 5 min. Protein bands were detected using a ChemiDoc imaging system (Bio-Rad).

### Confocal microscopy and image analysis

Images were acquired using either a Nikon A1R+HD confocal on a TiE inverted microscope (Nikon Instruments Inc., Japan, Nikon NIS-Elements imaging analysis software) using 60x/1.4 NA oil-immersion objective or on a Leica Stellaris 8 tau-STED Super Resolution Microscope (Leica Microsystems, Germany, LAS X software) using 93x/1.30 NA glycerol-immersion objective. All images were acquired at a frame size of 1,024 pixels. Optical Z-stacks were collected using the optimal Z-step interval calculated by the image acquisition software. Control and AP2-L-KO samples were acquired using identical settings, including laser power, detector gain, pinhole size, pixel dimensions, and Z-step interval. Images were processed and analyzed using Fiji/ImageJ (version 2.1.0 with Java 1.8.0) and Imaris (version 9.7; Bitplane, Oxford Instruments). Imaris was used for 3D image rendering, surface generation, object detection, and quantitative measurements. Within each experiment, identical image-processing and object-segmentation parameters were applied to Control and AP2-L-KO samples.

### Quantification of LC3 and ApoH association

Confocal Z-stack images were imported into Imaris (version 9.7). A surface representing the parasite was generated using UIS4 staining. A separate surface representing the host-cell nucleus was generated from the Hoechst signal and used as a spatial reference for generating surfaces corresponding to LC3 or ApoH fluorescence within the host cell. To quantify LC3 or ApoH associated with the parasite, a distance transformation was applied to the parasite surface to generate a region extending 2 µm beyond the PVM surface. A new surface was then created from the LC3 or ApoH fluorescence signal overlapping these 2 µm regions. For each EEF, parasite volume and the fluorescence intensity mean and sum of LC3 or ApoH within the 2 µm shell surrounding the PVM were obtained for subsequent analysis.

### RNA isolation from SG sporozoites and Sequencing

Total RNA was isolated from SG sporozoites using the RNeasy Mini Kit (Qiagen) according to the manufacturer’s instructions and immediately stored at −80°C. RNA concentration and purity were determined using Thermo NanoDrop One-C (Thermo Scientific). RNA integrity was evaluated using an Agilent Bioanalyzer. The assessed samples exhibited variable RNA integrity, with RNA integrity number values ranging from 4 to 7. Strand-specific mRNA libraries were prepared using the TruSeq Stranded mRNA Sample Preparation Kit LS (Illumina) according to the manufacturer’s instructions. Approximately 100-500 ng of total RNA was used as an input for each sample. Polyadenylated RNA was enriched using oligo-dT-coated magnetic beads, followed by first-strand cDNA synthesis using random primers. During second-strand synthesis, dUTP was incorporated in place of dTTP to preserve library strand specificity. Following end repair and adaptor ligation, libraries were amplified using 15 cycles of PCR. Final libraries were quantified using a Qubit 3.0 fluorometer (Thermo Fisher Scientific), and fragment-size distributions were assessed using an Agilent Bioanalyzer. Libraries were pooled at equimolar concentrations and sequenced by Novogene, Singapore, on an Illumina NovaSeq X platform to generate 150 bp paired end reads.

### RNA-sequencing data processing and differential expression analysis

RNA-sequencing data were processed using the Galaxy platform (galaxyproject.org). Sequencing read quality was assessed using FastQC (version 0.12.1). Adaptor sequences and low-quality bases were removed using Trimmomatic (version 0.39+galaxy2). Sliding-window trimming was performed over five bases and reads were trimmed when the mean Phred quality score fell below 20. The *P. berghei* ANKA reference genome and corresponding gene annotation were obtained from PlasmoDB (release 68). Trimmed paired-end reads were aligned to the reference genome using HISAT2 (version 2.2.1+galaxy1). Alignment and scoring parameters were maintained at their default settings. Gene-level read counts were generated using HTSeq-count (version 2.0.9+galaxy0). Reads were counted in union mode using exon features. Differential expression analysis of AP2-L KO and Control sporozoites incubated was performed using DESeq2 (version 2.11.40.8+galaxy0). Genes with an adjusted P-value below 0.05 and an absolute log_2_ fold change greater than 1 were considered significantly differentially expressed. Gene Ontology enrichment analysis was performed using the GO Enrichment tool in PlasmoDB. Upregulated and downregulated DEGs were analyzed separately. Enrichment was evaluated for biological process, molecular function, and cellular component ontologies using all annotated *P. berghei* genes as the background set. Enrichment significance was determined using Fisher’s exact test and GO terms with P-value below 0.05 were reported as enriched. De novo motif-enrichment analysis was performed using HOMER (version 5.1;). Genomic regions from DEGs extending from 1000 bp upstream to 300 bp downstream of the transcription start site were used were supplied as BED intervals and analyzed against the *P. berghei* ANKA reference genome using the -size given option. HOMER-generated genomic sequences served as the background.

### Statistical analysis

All statistical tests were performed in GraphPad Prism 10.6. P values <0.05 were considered statistically significant. Statistical details, including number of independent experiments, statistical tests used and definition of error bars, are provided in the figure legends.

## Data Availability

RNA-seq reads generated in this study have been deposited in the National Center for Biotechnology Information (NCBI) Sequence Read Archive (SRA) and are accessible through BioProject ID: PRJNA1551474.

## Acknowledgements

We acknowledge support from the National Institute of Health high-end instrumentation grant S10OD25182, which supported acquisition of the Leica Stellaris 8 tau-STED system used in this study.

## Supplementary Table(s)

Supplementary Table 1. Provided as a separate Excel file

**Supplementary Table 2.** Prepatent time when tested on immunodeficient mice.

| Mice | Number of injected sporozoites | Number of infected mice | Number of patent mice | Prepatent time (days) |
| --- | --- | --- | --- | --- |
| Control (C57BL/6) | 5 x 10<sup>3</sup> | 9 | 0 | No infection |
| IFNR -/- | 5 x 10<sup>3</sup> | 8 | 0 | No infection |

**Supplementary Table 3.**
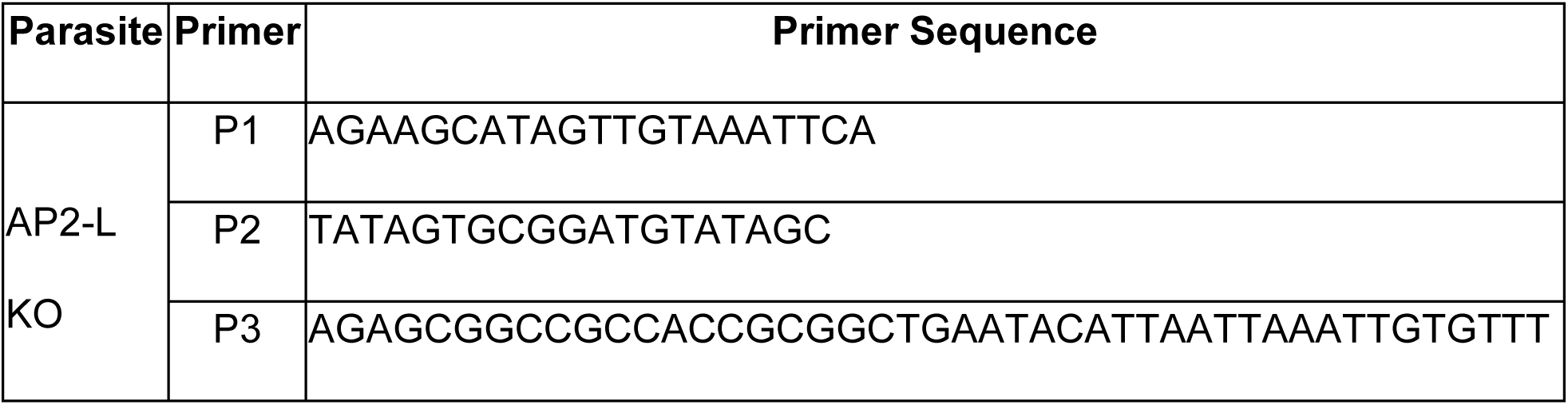
Primers used in this study.

**Supplementary Table 4.** Information regarding the primary antibodies.

| **Antibody** | **Host species** | **Supplier/Catalog number** | **Dilution** |
| --- | --- | --- | --- |
| Anti-APOH | Rabbit<br>Polyclonal | Sigma; HPA003732 | 1:250 |
| Anti-COPB2 | Rabbit<br>Polyclonal | Proteintech; 14733-1-AP | 1:200 |
| Anti-CSP (3D11) | Mouse<br>Monoclonal | Gift from Dr. Victor Nussenzweig | 1µg/mL |
| Anti-EXP1 | Chicken<br>Polyclonal | Gift from Dr. Volker Heussler | 1:500 |
| Anti-Gamma tubulin | Rabbit<br>Polyclonal | Invitrogen; PA534815 | 1:1000 |
| Anti-GFP (212) | Rabbit<br>Polyclonal | Gift from Dr. Conrad Beckers | 1:1000 |
| Anti-GM130 | Mouse<br>Monoclonal | BD Transduction Laboratories; 610822 | 1:500 |
| Anti-Golgin-97 (D8P2K) | Rabbit<br>Monoclonal | Cell Signaling Technology; 13192T | 1:100 |
| Anti-HSP70 (4E4) | Mouse<br>Monoclonal | Kind gift from Dr. Moriya Tsuji | 2µg/mL |
| Anti-LC3 (MAP1LC3A/B) | Rabbit<br>Polyclonal | BioRad; AHP2167T | 1:100 |
| Anti-UIS4 | Goat<br>Polyclonal | SicGen; A0042-200 | 1:1000 |
| Alexa Fluor 488 anti-rabbit | Donkey<br>Polyclonal | Invitrogen; Cat # A-21206 | 1:3000 |
| Alexa Fluor 594 anti-rabbit | Donkey<br>Polyclonal | Invitrogen; A21207 | 1:3000 |
| Alexa Fluor 488 anti-chicken | Goat<br>Polyclonal | Invitrogen; A11039 | 1:3000 |
| Alexa Fluor 488 anti-goat | Donkey<br>Polyclonal | Invitrogen; A11055 | 1:3000 |
| Alexa Fluor 594 anti-goat | Donkey<br>Polyclonal | Invitrogen; A-11058 | 1:3000 |
| Alexa Fluor 488 (F(ab')<sub>2</sub>) anti-mouse | Goat<br>Polyclonal | Invitrogen; A11017 | 1:3000 |

## Supplementary Figures

**Supplementary Figure 1.**
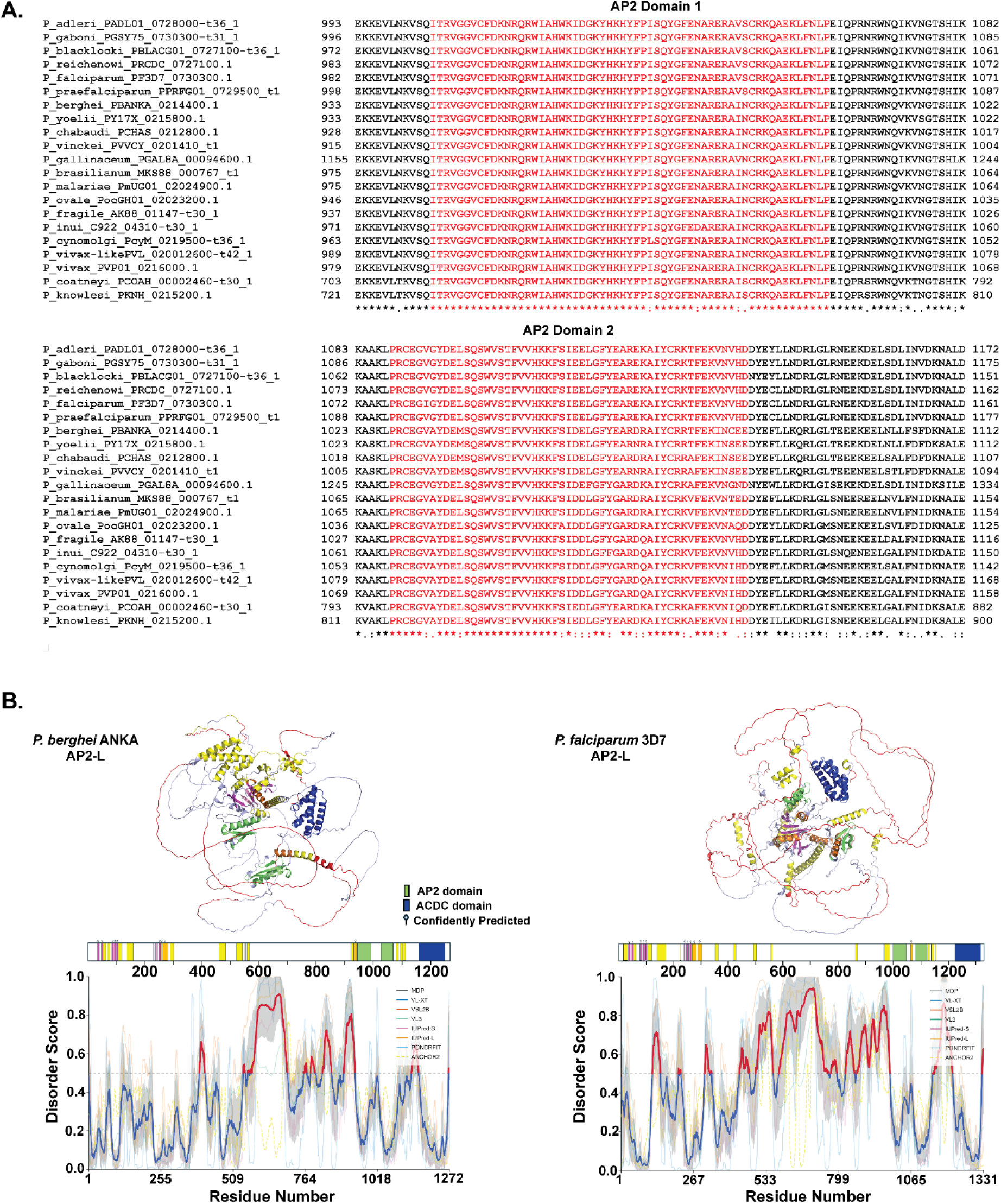
Sequence conservation, AlphaFold 3 structural predictions, and intrinsic disorder profiles of AP2-L orthologs. **(A)** Multiple sequence alignment of regions encompassing the first and second AP2 domains from 21 *Plasmodium* AP2-L orthologs. Numbers indicate amino acid positions within the corresponding full-length proteins. Asterisks denote residues identical across all sequences; colons and periods indicate residues with strongly and weakly similar physicochemical properties, respectively. **(B)** AlphaFold 3 structural model predictions of the full-length AP2-L from *P. berghei*, and *P. falciparum*. The AP2 domains, ACDC domain, α-helical regions, and β-strand regions are colored as indicated. Residues predicted to be disordered by Rapid Intrinsic Disorder Analysis Online (RIDAO) are highlighted in red in the structural models. A linear schematic beneath each model indicate the positions of the domains and secondary-structure elements. Corresponding RIDAO disorder profiles are shown below each schematic and aligned by amino acid positions.

**Supplementary Figure 2.**
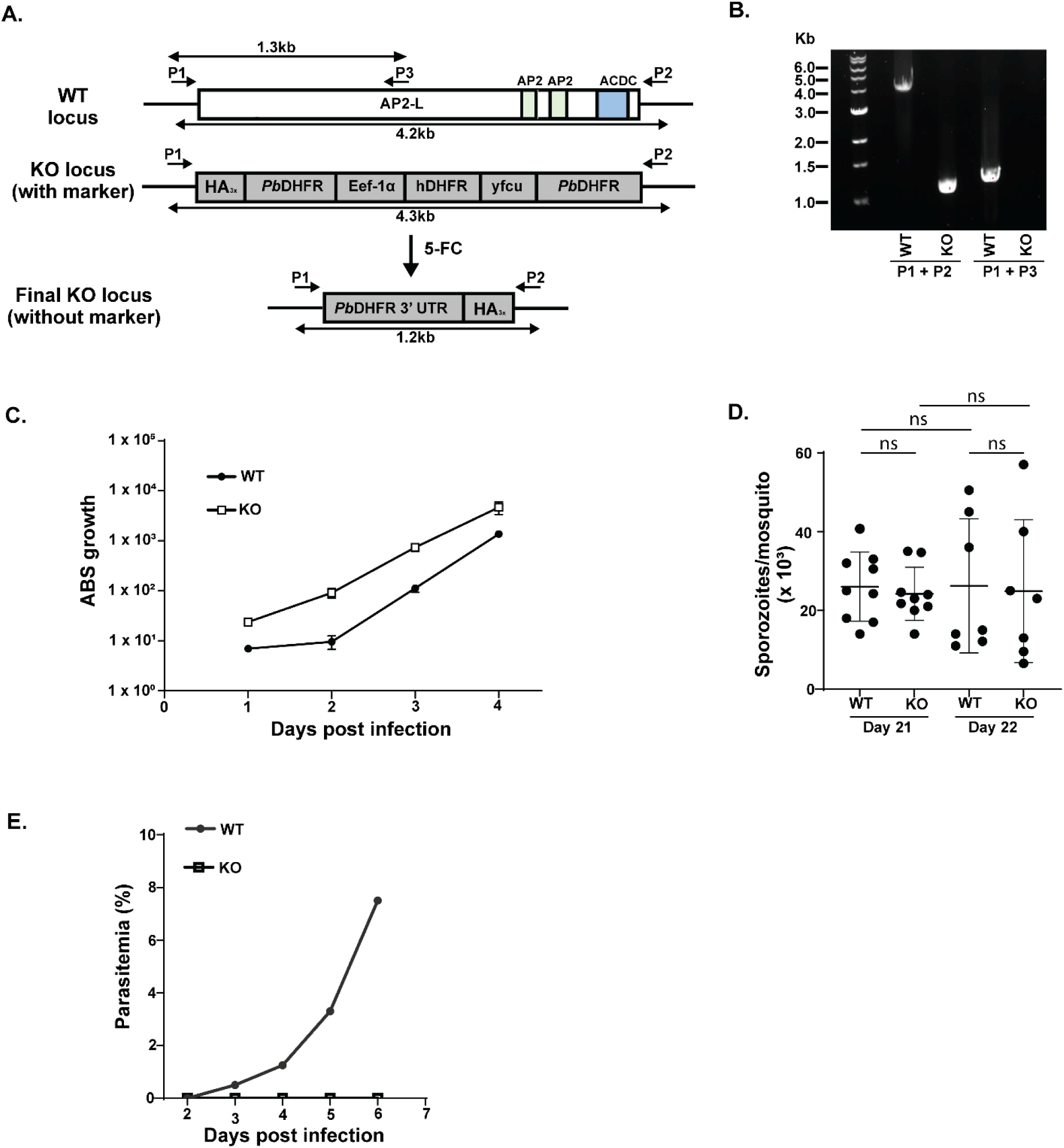
Generation of *P. berghei* AP2-L KO parasites. **(A)** Schematic showing generation of marker free mutant parasite through modification of the ap2-l locus **(B)** PCR confirmation of the modification of the ap2-l locus. Primer positions are indicated in panel A. **(C)** Growth rate of AP2-L KO asexual blood stages. **(D)** Infectivity of AP2-L KO sporozoites in *A. stephensi* mosquitoes. Salivary gland sporozoites were collected on day 21 and 22 post blood meal. The average number of sporozoites per mosquito (×10^3^) was calculated. **(E)** Growth rate of AP2-L KO asexual blood stages after sporozoite infection of mice via mosquito bite. Data represent the mean ± SD from 3 or more independent experiments. Statistical significance was determined using an unpaired two-tailed Student’s t-test (ns, P > 0.05).

**Supplementary Figure 3.**
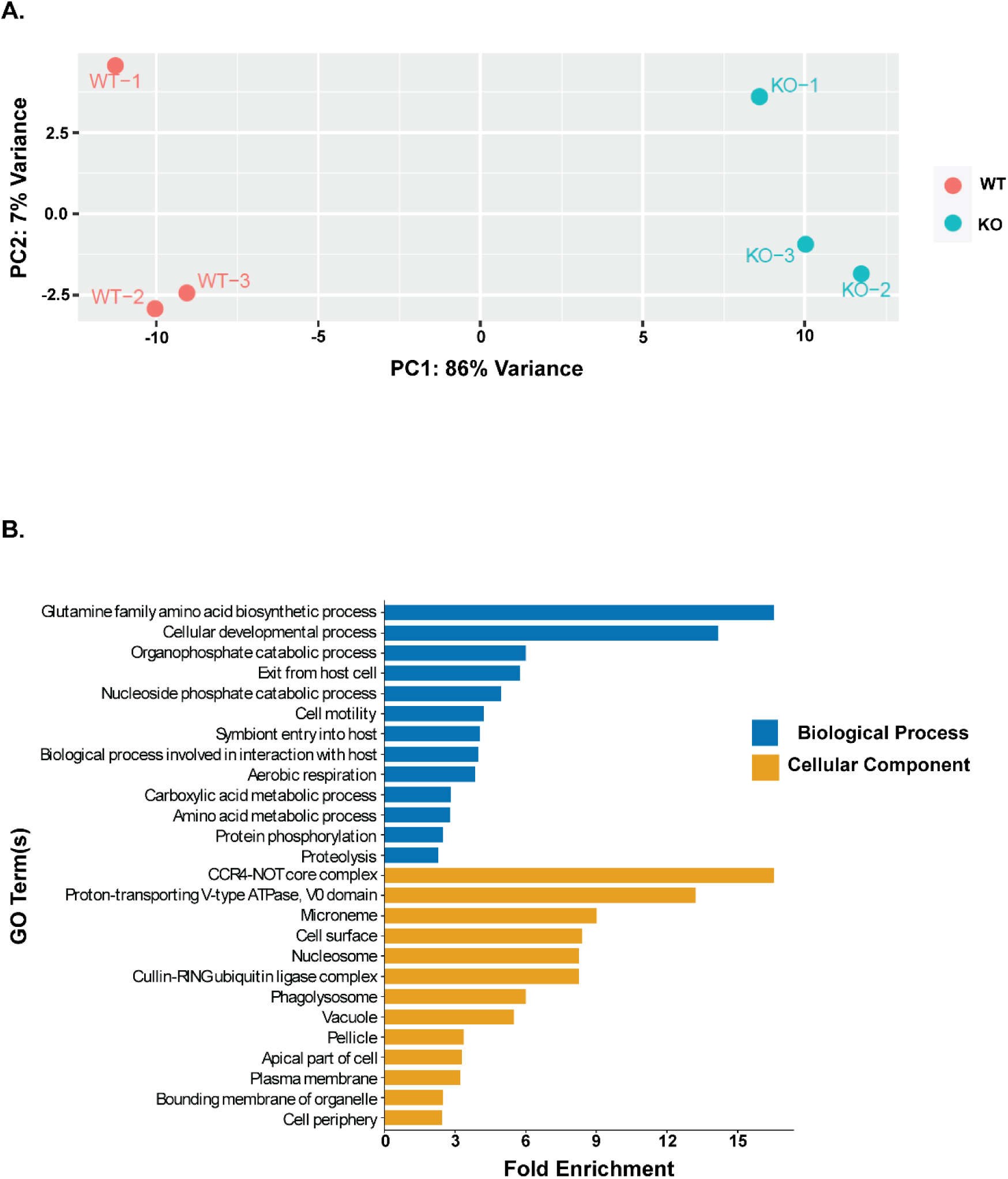
**(A)** Principal component analysis (PCA) performed on normalized gene expression counts to assess global transcriptional differences between Control and AP2-L KO sporozoites. Each point represents a single biological replicate, with Control and AP2-L KO groups indicated by different colors. PC1 explains 86% of the variance while PC2 explains 7% of the variance. **(B)** Gene Ontology (GO) enrichment analysis of low abundance genes in AP2-L KO where the CATGCA motif was identified.

**Supplementary Figure 4.**
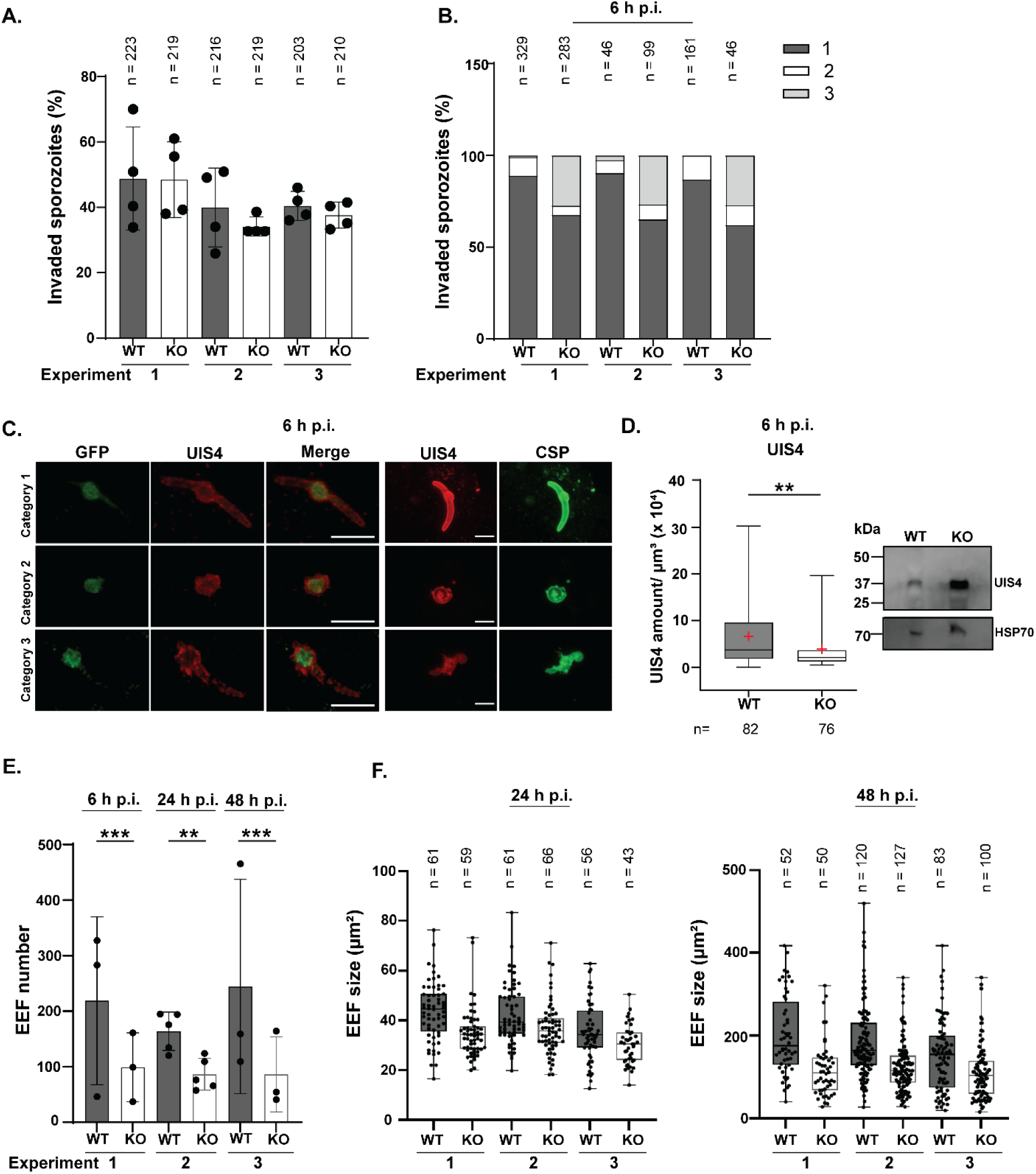
AP2-L is required for sporozoite transformation and liver-stage growth. **(A)** Invasion of HepG2 cells by Control and AP2-L KO sporozoites was quantified using double differential immunofluorescence staining. Invasion efficiency was calculated as the percentage of intracellular sporozoites relative to the total number of sporozoites. Shown are results from three independent experiments. **(B)** Sporozoite transformation into EEFs was quantified at 6 h p.i. using GFP to visualize the parasite cytoplasm (green) and UIS4 to visualize the PVM (red). Parasites were assigned to one of three categories based on parasite morphology and UIS4 distribution. Shown are results from three independent experiments. **(C)** Representative immunofluorescence images of the three categories are shown. Scale bars, 10 µm. Corresponding STED images show UIS4 and CSP localization in each category. Scale bars, 3 µm. **(D)** UIS4 fluorescence signal surrounding each GFP-positive sporozoite was measured within a 2 µm region and normalized to the GFP-defined sporozoite volume. Adjacent immunoblot shows UIS4 abundance in Control and AP2-L-KO SG sporozoites, with HSP70 used as a loading control. **(E)** Results from three independent experiments to quantify EEF numbers at 6, 24, and 48 h p.i.. Statistical significance was determined using an unpaired two-tailed Student’s t-test (**P ≤ 0.01; ***P ≤ 0.001) **(F)**. Results from three independent experiments to quantify EEF size at 24 and 48 h p.i.

**Supplementary Figure 5.**
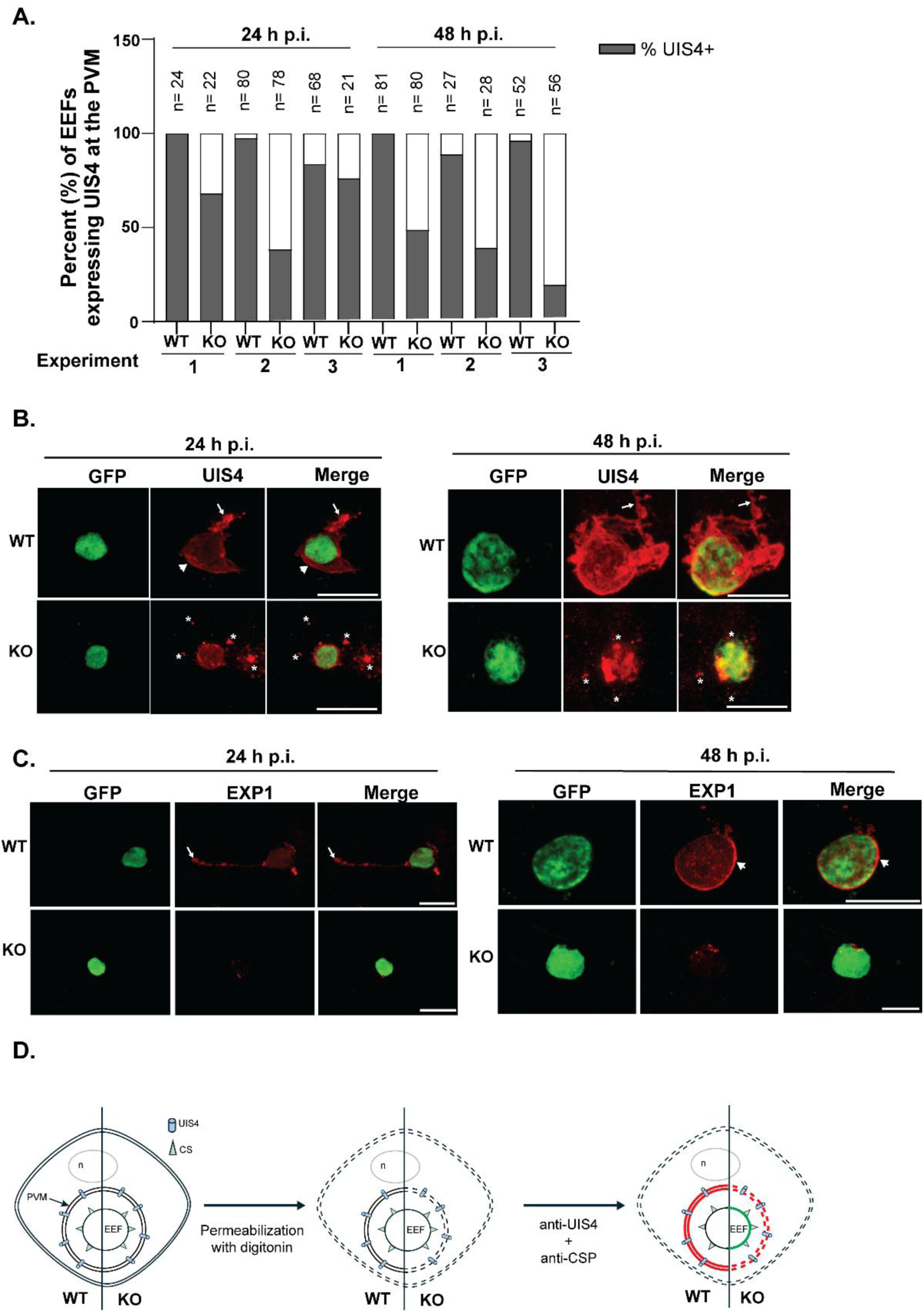
Loss of AP2-L disrupts localization of PVM proteins and compromises PVM integrity (A). Results from three independent experiments quantifying the percentage of Control and AP2-L-KO displaying UIS4 localization at the PVM at 24 and 48 h p.i. **(B)** Representative immunofluorescence images of infected HepG2 cells at 24 or 48 h p.i. Parasites were stained for GFP to visualize the parasite cytoplasm and for UIS4 to visualize the PVM. Arrows indicate PVM extensions, arrowheads indicate the PVM surrounding the parasite, and asterisks indicate UIS4-positive puncta. Scale bars, 10 µm. **(C)** Representative immunofluorescence images of infected HepG2 cells at 24 and 48 h p.i. Parasites were stained for GFP and the PVM protein EXP1. Arrows indicate PVM extensions and arrowheads indicate the PVM surrounding the parasite body. Scale bars, 10 µm. **(D)** Schematic of the digitonin-permeabilization assay used to evaluate PVM integrity.

**Supplementary Figure 6.**
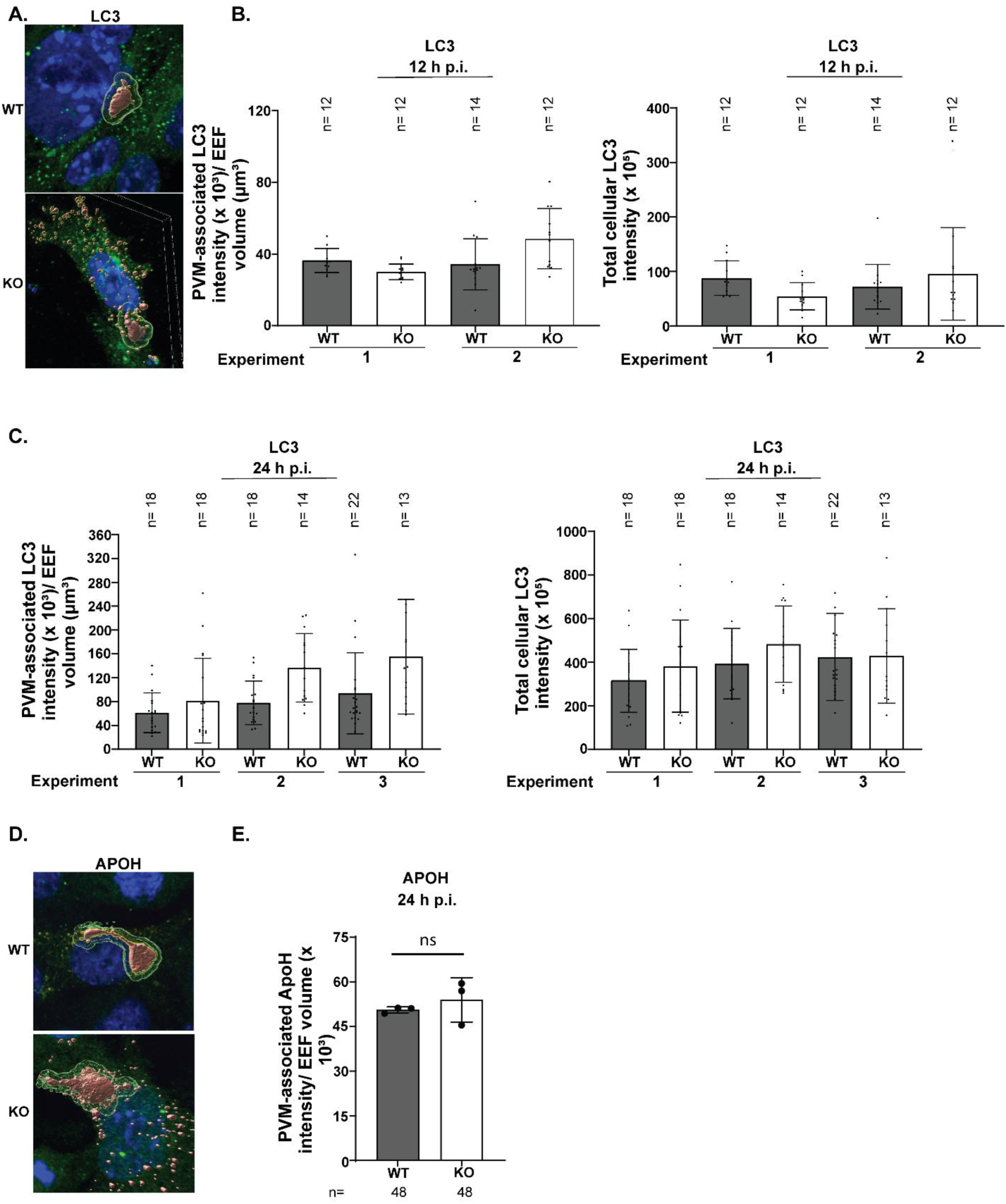
Quantification of LC3 and ApoH surrounding Control and AP2-L-KO EEFs. **(A)** Representative Imaris reconstructions illustrating the measurement of LC3 surrounding EEFs. The EEF is displayed as an opaque red surface and LC3 fluorescence within a 2 µm region surrounding the EEF is displayed in transparent green. **(B)** PVM-associated LC3 fluorescence intensity normalized to EEF volume and total cellular LC3 fluorescence intensity at 12 h p.i. **(C)** PVM-associated LC3 fluorescence intensity normalized to EEF volume and total cellular LC3 fluorescence intensity at 24 h p.i. **(D)** Representative Imaris reconstructions illustrating the measurement of ApoH surrounding Control and AP2-L KO EEFs. The EEF is displayed as an opaque red surface, and ApoH fluorescence within a 2 µm region surrounding the EEF is displayed in transparent green. **(E)** PVM-associated ApoH fluorescence intensity normalized to EEF volume at 24 h p.i. Statistical significance was determined using an unpaired two-tailed Student’s t-test (ns, P > 0.05).

**Supplementary Figure 7.**
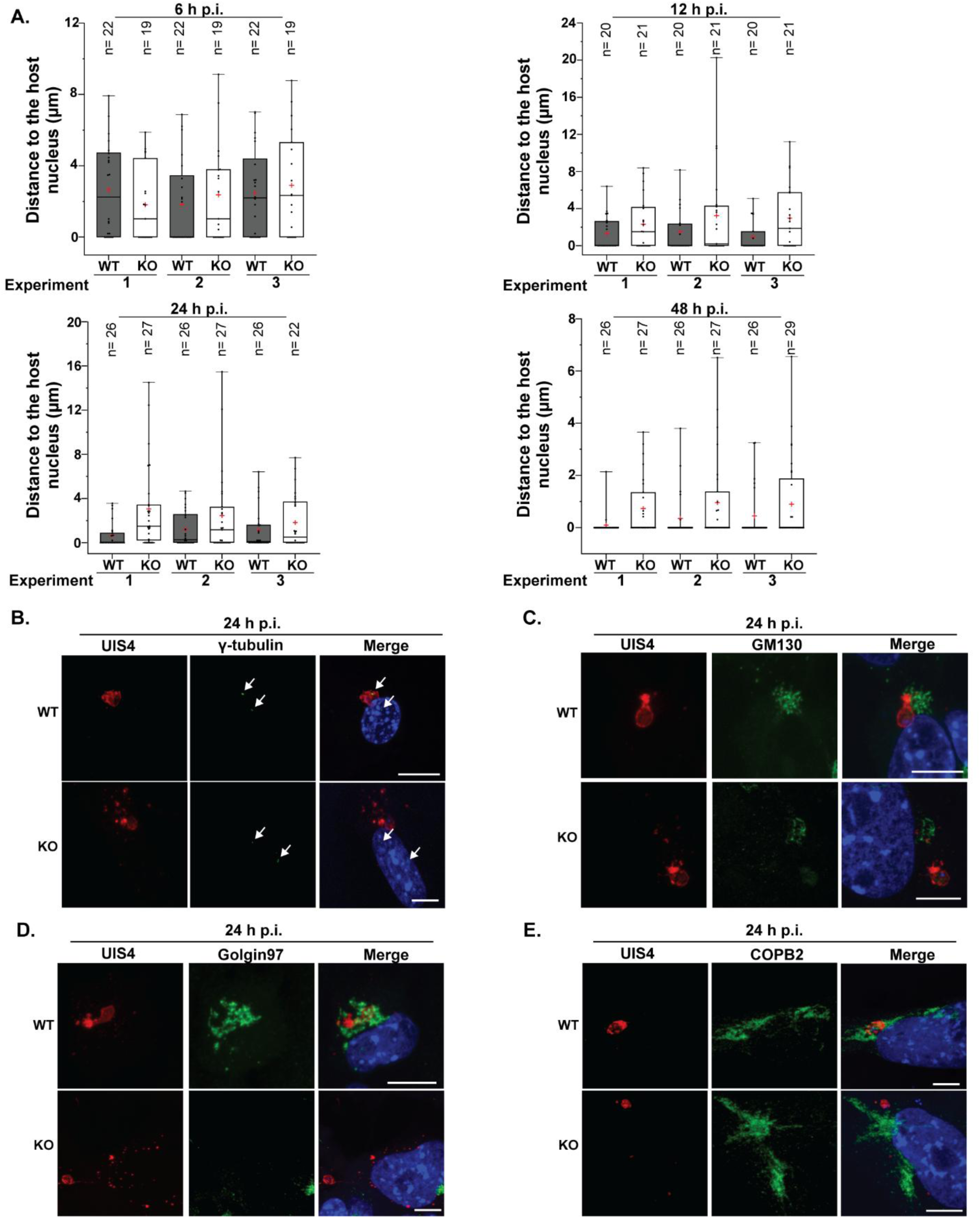
Loss of AP2-L alters EEF perinuclear positioning, reduces association with host γ-tubulin foci and with host vesicular pathways. **(A)** Distance of Control and AP2-L KO EEFs from the host-cell nucleus at 6, 12, 24, and 48 h p.i. The distance between the UIS4-positive PVM and the periphery of the host-cell nucleus was measured from maximum-intensity projections. Results from three independent experiments are shown for each time point. **(B)** Representative immunofluorescence images of Control- and AP2-L KO infected HepG2 cells at 24 h p.i. showing UIS4-positive EEFs (red), γ-tubulin (green), and Hoechst-labeled nuclei (blue) in the merged images. White arrows indicate γ-tubulin foci. **(C)** Representative images showing UIS4 (red), the cis-Golgi marker, GM130 (green) and Hoechst-labeled nuclei (blue) in the merged images at 24 h p.i. **(D)** Representative images showing UIS4 (red), the trans-Golgi marker, Golgin-97 (green) and Hoechst-labeled nuclei (blue) in the merged images at 24 h p.i. **(E)** Representative images showing UIS4 (red), COPB2 (green), and Hoechst-labeled nuclei (blue) in the merged images at 24 h p.i.

